# Microsaccade Selectivity as Discriminative Feature for Object Decoding

**DOI:** 10.1101/2024.04.13.589338

**Authors:** Salar Nouri, Amirali Soltani Tehrani, Niloufar Faridani, Ramin Toosi, Jalaledin Noroozi, Mohammad-Reza A. Dehaqani

## Abstract

Microsaccades, a form of fixational eye movements, maintain visual stability during stationary observations. Previous studies have provided valuable insights into the relationship between microsaccade characteristics and external stimuli. However, the dynamic nature of microsaccades provides an opportunity to explore the mechanisms of information processing, particularly object decoding. This study examines the modulation of microsaccadic rates by different stimulus categories. Our experimental approach involves an analysis of microsaccade characteristics in monkeys and human subjects engaged in a passive viewing task. The stimulus categories comprised four primary categories: human, animal, natural, and man-made. We identified distinct microsaccade patterns across different stimulus categories, successfully decoding the stimulus category based on the microsaccade rate post-stimulus distribution. Our experiments demonstrate that stimulus categories can be classified with an average accuracy and recall of up to 85%. Our study found that microsaccade rates are independent of pupil size changes. Neural data showed that category classification in the inferior temporal (IT) cortex peaks earlier than microsaccade rates, suggesting a feedback mechanism from the IT cortex that influences eye movements after stimulus discrimination. These results exhibit potential for advancing neurobiological models, developing more effective human-machine interfaces, optimizing visual stimuli in experimental designs, and expanding our understanding of the capability of microsaccades as a feature for object decoding.

## Introduction

The human visual system operates a remarkable interaction between visual exploration and the brain’s neural mechanisms (Titchener et al., 2020). Within this system, eye movements, characterized by their frequent shifts in gaze, constitute fundamental and recurrent movements pivotal in facilitating our visual exploration of the environment (Veneri et al., 2011). In addition, within the neural system, these movements contribute to interpreting and categorizing visual stimuli during object classification (Stacchi et al., 2019).

Different types of eye movements have different roles in visual perception. Saccades, rapid eye movements, and shift gaze between points of interest contribute to quick visual exploration and attention. Microsaccades, referred to as fixational eye movements, are small involuntary motions during fixation, helping prevent retinal adaptation to stationary stimuli (Mellor & Psouni, 2021). They aid in continuously sampling our visual environment, thus enhancing our interpretation of the environment. This assistance in visual scene interpretation enables a deeper understanding of the environment, recognizing faces, and supporting various visually demanding tasks (Chen et al., 2022; Alexiev & Vakarelsky, 2022). Meanwhile, smooth pursuit movements enable the eyes to follow moving objects smoothly, ensuring consistent and precise visual tracking (Barnes, 2008; Souto & Kerzel, 2021; Parisot et al., 2021). These diverse eye movements collectively support the brain’s ability to recognize objects effectively.

The distinction between saccades and microsaccades primarily centers on their amplitude, duration, peak velocity, and frequency. Saccades have larger amplitudes, typically ranging from more than 1.0° to several degrees, occurring multiple times per second (more than 2-3 Hz). They typically last around 20 to 200 milliseconds and reach maximum velocity greater than 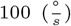 (Land, 2012; Otero-Millan et al., 2008; Gibaldi & Sabatini, 2021).

In contrast, microsaccades exhibit significantly smaller amplitudes, peak velocities, and durations (Otero-Millan et al., 2008; Gibaldi & Sabatini, 2021), unfold at a slower rate, typically occurring only a few times per second (1 Hz, (Costela et al., 2014)). Microsaccade amplitude, generally defined as saccades smaller than 1.0° of visual angle (Martinez-Conde et al., 2013; Melloni et al., 2009), could also be influenced by the size of the stimulus and the fixation target (Otero-Millan et al., 2013; McCamy et al., 2013). Microsaccades often attend the direction of visual spatial attention, increasing nerve cell activity at the focused spot during visual fixation. Despite their small size, microsaccades hold an essential role in visual perception by stabilizing the image on the retina, ensuring a consistent and clear view of our surroundings (Kasneci et al., 2022). They also contribute to efficient visual representations of the environment by adapting to local regularities induced by small spatial displacements over time (Rolfs et al., 2007). One contributing mechanism often associated with the generation of microsaccades involves the ongoing effort of the oculomotor system to sustain fixation on a single point despite subtle variations in stimuli or background noise. Tiny fluctuations can cause our eyes to drift away from the target, resulting in blurred vision when fixation on an object. Our oculomotor system generates microsaccades to bring the object back onto the fovea to maintain visual acuity (Peel et al., 2016).

Microsaccades can change brain processing and are subject to the influence of various cognitive factors, including task difficulty and attention (Engbert & Kliegl, 2003; Siegenthaler et al., 2014; Engbert, 2006). The microsaccade rate, termed the microsaccade signature, represents the frequency of microsaccades occurring within specific time intervals, contributing to regulating visual perception. The occurrence of microsaccade rate has been demonstrated to be influenced by various factors, including task demands and attentional processes (Kim et al., 2015). Moreover, focusing on a particular location increases the microsaccade rate, facilitating obtaining additional information about the focused stimulus through guiding gaze shifts. Conversely, without relevant stimuli requiring attention, the microsaccade rate decreases (Thielen et al., 2019). Furthermore, the microsaccade rate reduces temporal attention directed at visual targets and increases attentional load across both visual and non-visual tasks (Kasneci et al., 2022). In addition, when participants view complex or dynamic stimuli, such as naturalistic scenes, there is an increase in the microsaccade rate compared to when they view simple or static stimuli like patterns (Brych et al., 2021). Among the factors influencing microsaccade rate, an important variable is the stimulus category (Hafed & Ignashchenkova, 2013). Numerous studies have examined the impact of stimulus categories on microsaccade rates during visual perception tasks. The stimulus category contains the characteristics of visual stimuli presented to participants during experimental tasks. The stimuli are categorized based on various criteria: complexity, brightness, contrast, color, and spatial frequency content (Essig et al., 2020). Siegenthaler et al. (2014) found that microsaccade rates were higher during naturalistic scene perception compared to the perception of simple patterns.

Similarly, Engbert & Kliegl (2003) reported increased microsaccade rates with increasing stimulus complexity during visual discrimination tasks. In addition, following a visual stimulus onset, the microsaccade rate displays a distinctive pattern of initial inhibition before reaching a new peak approximately 200–400 milliseconds after the stimulus onset (Engbert & Kliegl, 2003; Turatto et al., 2007). The high-level properties of the stimulus impact the peak in saccade rate after initiating a visual stimulus. This increase in the saccade rate tends to be higher for objects than non-object stimuli, aligning with the concept of the microsaccade role in object recognition (Yuval-Greenberg et al., 2008; Hassler et al., 2011; Keren et al., 2010). Subsequently, microsaccades have been associated with the visual detection of dim objects, although their impact on peripheral versus central vision remains unclear. The ability of microsaccades to recover small objects within the foveal region, however, is still a subject of debate because the evaluated target size extended beyond the foveal boundaries (Zhang et al., 2020; Toscani et al., 2017). It is suggested that task and stimulus categories can modulate microsaccade rates, with complex stimuli requiring heightened attention and cognitive processing, thus resulting in an increased microsaccade rate (Intoy & Rucci, 2020; Di Stasi et al., 2016).

Despite the recognized importance of microsaccades in visual information processing, the study of fixational eye movements has yet to receive sufficient attention in research. The impact of different stimulus categories on microsaccade rates represents an area requiring more exploration. Although these categories may affect microsaccade rates in distinct or similar ways following stimulus onset, a detailed examination of their effects remains sparse. Bridging this research gap is essential for understanding microsaccade selectivity in object decoding. This necessitates a focused study to test the hypothesis that different stimulus categories uniquely modulate microsaccade rates, influencing their selectivity in decoding objects. To explore this hypothesis, we have designed an experiment to assess the contribution of microsaccades to decoding object categories.

Our study focused on exploring the capacity of microsaccade rates as distinguishing features for recognizing objects, aiming to improve efficiency and deepen our understanding of behavioral aspects related to object decoding. We conducted an analysis focusing on the relationship between microsaccade patterns and object recognition. This analysis used data obtained from two rhesus monkeys and twenty-five human participants. The study contained observations across four primary stimulus categories: human, animal, natural, and man-made. We measured and analyzed microsaccade rates using a rapid serial visual presentation (RSVP) protocol. Our findings highlighted distinct average microsaccade rates among categories, with the animal category exhibiting the highest rates and the man-made category displaying the lowest. Employing a support vector machine (SVM), we utilized microsaccade rate distributions to predict stimulus categories, achieving high accuracy and recall rates between 70% and 85% in post-stimulus distribution. We also conducted additional analyses of pupil size and neural data to explore the underlying factors influencing microsaccade rates. This analysis revealed that pupil size did not significantly differ across different stimulus categories for both human and monkey datasets. Hence, the observed modulation of microsaccades is not directly related to changes in pupil size. Furthermore, our neural data analysis indicated that the peak accuracy for classifying stimulus categories in the inferotemporal (IT) cortex occurs earlier than the peak microsaccade rate. So, it suggests a potential feedback mechanism from the IT cortex influencing eye movements after stimulus discrimination rather than a direct relationship with brain state changes such as those indicated by pupil size. This study enhances our understanding of microsaccades as pivotal tools for decoding stimulus categories based on their rates and emphasizes their potential role in object recognition.

## Methods

### Participants

Our study utilized two distinct datasets obtained from separate experiments to explore microsaccade rates across various stimulus categories.

#### Experiment 1: Animal Experiment

In this study, two mature male rhesus monkeys were employed for the first dataset. Both tests were conducted at the School of Cognitive Science at Research in Fundamental Sciences (IPM). The Society for Neuroscience Guidelines and Policies and the National Institutes of Health Guide for the Care and Use of Laboratory Animals were followed during every experiment step. The Council for the Institute of Fundamental Science approved all surgical, behavioral, and experimental operations protocols. At the start of every trail, the monkeys’ initial eye locations were fixed on the center of the monitor.

#### Experiment 2: Human Participants

The second dataset was collected from 25 human participants, including 15 men and 10 women. They were equipped with head-mounted eye trackers featuring an infrared radiation (IR) camera to capture their eye movements. Detailed consent forms outlining the study’s purpose, procedures, potential risks, and confidentiality measures were provided to all human participants. This process ensured that consent was fully informed and voluntarily given, documented in writing before participation.

### Stimuli

Our study utilized 155 grayscale images from the Hemera Photo Items commercial picture library. These images covered different subjects, including humans, animals, natural scenes, and man-made objects. To ensure consistency across the study, we exclusively selected images that depicted single objects in isolation. Visual representations of six examples of stimulus categories, which include human (face and body), animal (face and body), natural, and man-made, are presented in Fig. 1(A). The categories of animals, humans, faces, and bodies are grouped as animate, while man-made and natural categories fall under inanimate. Fig. 1(F) illustrates a clear and structured flowchart of stimulus categories used in our study.

**Fig. 1.**
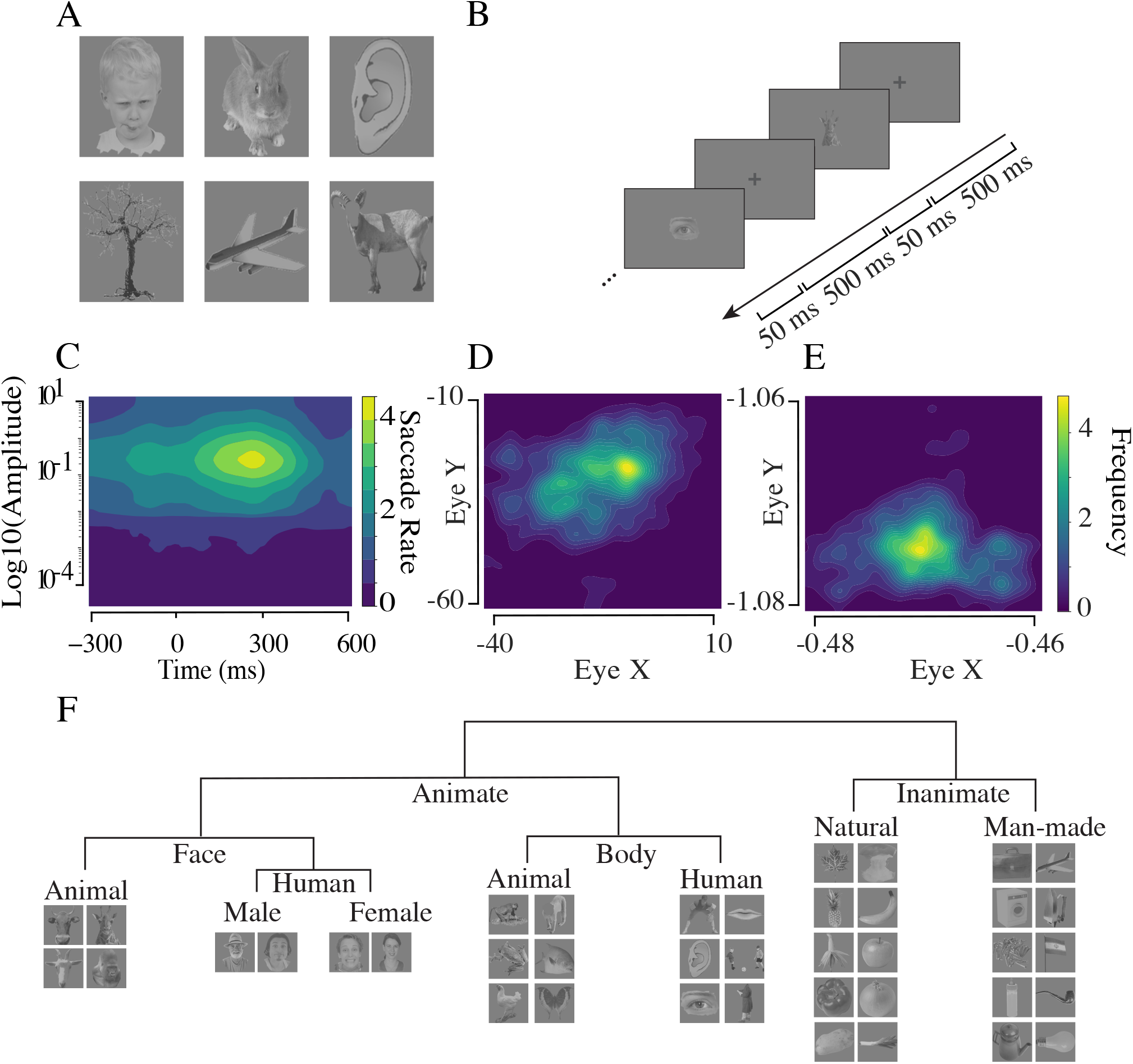
Experimental paradigm and eye movements. **A)** A sample of the stimulus Set: The stimulus set utilized in our experiment comprises grayscale images, including four main categories: humans, animals, man-made objects, and natural scenes. Human and animal stimuli are further categorized into faces and bodies, forming the animate category, while man-made and natural entities create the inanimate category. **B)** Experimental task Paradigm: Each trial involves participants focusing on a sequence of images while their eye movements are recorded. Employing the RSVP technique, every trial displayed 155 randomly chosen stimuli across various categories. The task sequence was initiated with a blank screen, succeeded by the presentation of a fixation point lasting 500 milliseconds. Subsequently, each stimulus appeared for a duration of 50 milliseconds. The participants’ eye positions were continually tracked and recorded throughout the trials. **C)** Distribution of saccades and amplitudes over time: The aggregated distribution of saccade rates and amplitudes is presented across time bins of 15 milliseconds. The amplitude axis is represented using a logarithmic scale with a log amplitude of 0.05. Time zero marks the stimulus onset. **D)** Eye movements distribution for a trial in monkey: This plot illustrates a distribution of eye movements recorded in a monkey participant while viewing a specific stimulus. The color intensity signifies the frequency of eye movement occurrences during the experiment, with brighter colors indicating higher frequencies of eye positions. **E)** Eye movements distribution for a trial for a human subject: It presents a distribution of eye movements recorded in a human participant while viewing a specific stimulus. **F)** A structured flowchart showing the superordinate and mid-level stimulus categories used in our study.

#### Visual Stimuli Presentation

The presentation of visual stimuli was standardized using a 19-inch Eizo FlexScan S1910 monitor, with a screen resolution of 1280 *×* 1024 pixels. The monitor was positioned at a 65.5 cm viewing distance from the subject’s eye. Stimuli were centrally displayed on the screen, each measuring 500 *×* 500 pixels against a neutral gray background, resulting in a visual angle of 10 degrees both horizontally and vertically.

### Eye Movement Tracking

#### Animals’ Eye Tracking

The animals were put in specially designed primate seats, with their heads immobilized and a juice delivery tube inserted into their mouths as incentives. An infrared optical eye tracking device (EyeLink 1000 Plus Eye Tracker, SR Research Ltd, Ottawa, CA) was used to monitor eye position at a frequency of 2 KHz, with a resolution of less than 0.01 RMS. The EyeLink 1000 Plus Camera (SR Research Ltd, Ottawa, CA) and EyeLink PM910 Illuminator Module were positioned in front of the monkey to record eye movements. An LED-lit 24-inch monitor with a resolution of 1920 *×* 1080 and a refresh rate of 144 Hz was used to display visual stimuli, and it was placed 65.5 cm in front of the animal’s eyes. The calibration procedure for the eye tracker involved a multi-point calibration method conducted at the start of each session to ensure accurate eye position tracking. Specifically, the monkeys were trained to fixate on a series of points displayed sequentially at known locations on the screen. These points typically cover the central and peripheral visual fields to provide a comprehensive calibration grid. The calibration process was repeated until the recorded eye positions matched the known positions of the calibration points with high accuracy. The primate seats were ergonomically designed to minimize movement. In overall, we collected 235 sessions with 1,246,379 trials for the monkey dataset.

#### Humans’ Eye Tracking

A remote monocular eye-tracker (Segal Tracker, Tehran, Iran) with a sampling frequency of 500 Hz and a spatial resolution of less than 0.2° RM was used to track the participant’s left eye. The stimulus presentation was controlled using the PsychToolBox (PTB) framework. The subject was positioned 55 centimeters in front of a 57 by 43.1 centimeter screen with a 1920 by 1080 pixel resolution to see the stimuli. As a result, the pixels on the screen had a viewing angle of 0.031 degrees. Before the experiment, a calibration process using a 5-point calibration grid was performed to ensure precise eye movement tracking. In overall, we gathered 74 sessions with 19,018 trials for human dataset.

### Task Paradigm

Our experimental procedure employed the RSVP method to explore the microsaccade rates across visual stimuli. Each session consisted of 5 blocks, with 155 stimuli randomly presented in each block. At the beginning of each block, a blank screen with a central fixation point was displayed for 500 milliseconds to establish a baseline for participants’ eye positions and standardize the starting condition for each trial. Following the baseline period, each stimulus was presented for 50 milliseconds. The stimulus included a variety of visual categories to examine the discriminative power of microsaccade rates. After each stimulus presentation, a blank screen was shown for an inter-stimulus interval ISI of 500 milliseconds. This interval was crucial for providing a clear separation between consecutive stimuli, allowing participants sufficient time to process each stimulus independently. Throughout the trials, participants’ eye positions were continuously tracked and recorded. For monkeys, the eye tracking was performed at a rate of 2000 Hz, while for humans, it was conducted at a rate of 500 Hz. Fig. 1(B) illustrates the experimental paradigm and presents examples from the stimulus set to understand better the task and the stimuli employed. The stimulus set, including their standardization, is the same for human and monkey experiments. The difference is only in the hardware setup. We used different hardware setups for monkeys and humans due to the specific requirements and constraints of each experimental setup:

### Eye Tracking Analysis

We employed a specific temporal range to analyze and detect eye movements, extending from 300 ms before the stimulus onset (baseline period) to 600 ms following the stimulus presentation (post-stimulus period). Any trial wherein a blink was detected during the temporal window or if the participant’s eye position was not accurately tracked throughout the specified range was excluded from subsequent analysis to have high-quality data. Following these criteria, approximately 3.2% of the collected data were considered unsuitable for analysis and thus excluded.

Saccade amplitude is a metric that quantifies the eye’s movement distance covered during a saccade. This can be measured in visual degrees (angular distance) or pixels, estimated by the Euclidean distance between consecutive fixation points (Holmqvist et al., 2011). Saccade velocity is a key indicator determined by analyzing the first temporal derivative of the gaze position data. It includes average saccadic velocity, an average speed over the entire duration of a saccade, and peak saccadic velocity, indicating the maximum speed achieved within a saccade (Holmqvist et al., 2011). Microsaccade duration is calculated by the interval from the onset of a microsaccade to its offset. Fig. 2 represents the saccade probability as a function of amplitude, duration, and peak velocity. Each detected saccade is analyzed to determine its amplitude, duration, and peak velocity. The y-axis, labeled saccade probability, indicates how frequently saccades with specific characteristics (amplitude, duration, peak velocity) occur within the dataset.

**Fig. 2.**
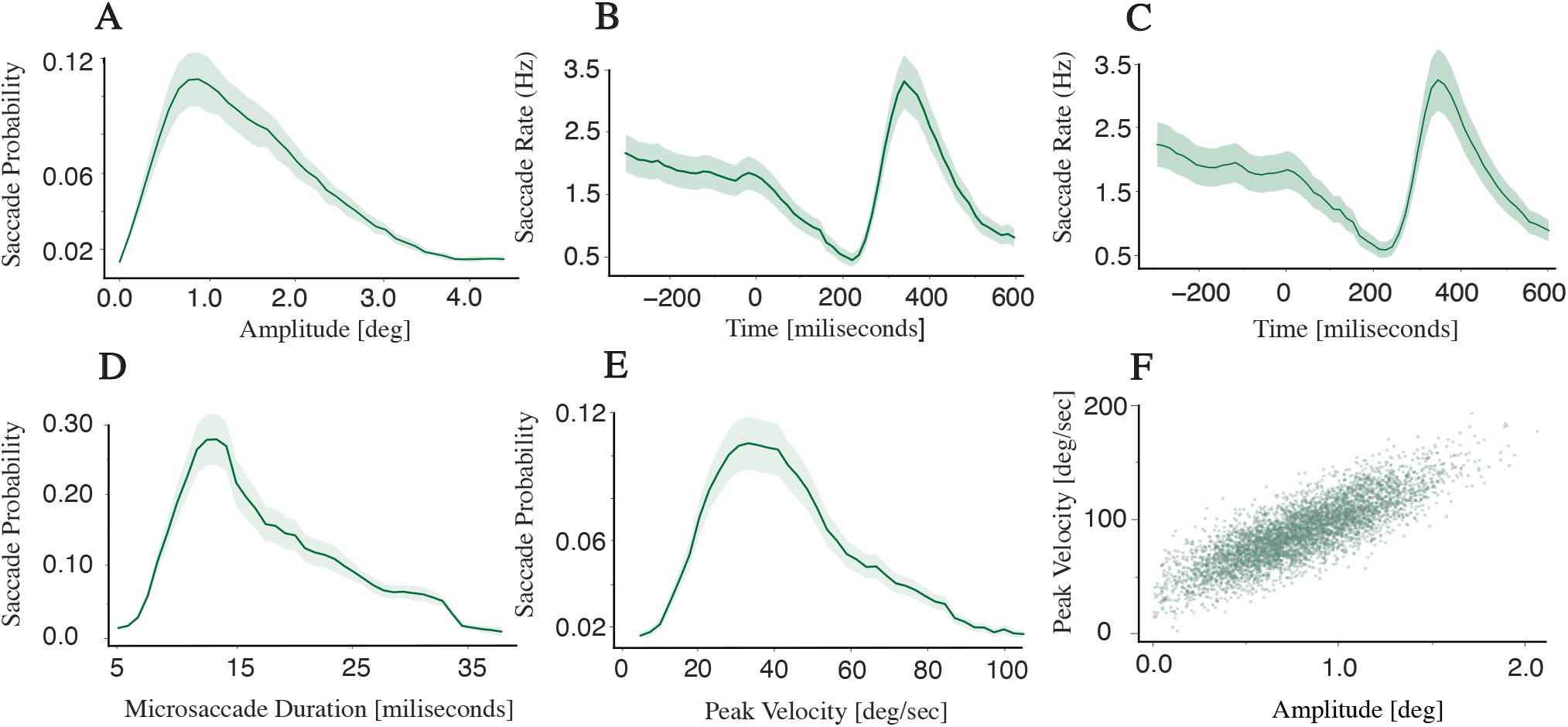
Validity of the detected saccades. **A)** Saccade amplitude distribution in monkeys: The distribution of saccade amplitudes observed in monkeys across all categories has a clear unimodal pattern, with the peak occurring at approximately 1.0 degrees of amplitude. **B)** Saccade rate over time in monkeys: The distribution of detected saccade rates in monkeys across all categories over time, with a bin width of 15 milliseconds, shows a distinct increase in the rate of microsaccades in 200–400 milliseconds following stimulus presentation. This distribution demonstrates an unimodal pattern after stimulus onset. **C)** Saccade rate over time in Humans: The distribution of saccade rate across all categories obtained from human data, with a bin width of 15 milliseconds, illustrates the similar distinct increase in the saccade rate around 200 - 400 milliseconds after stimulus onset. **D)** Saccade probability distribution by microsaccade duration: The distribution of saccade probability concerning microsaccade duration across all categories showcases an unimodal pattern, with the highest peak occurring at approximately 14 ms for microsaccade duration. Approximately 40% of the detected microsaccades are within the range of 10 to 25 ms duration. **E)** Saccade probability distribution by peak velocity: The distribution of saccade probability concerning the peak velocity of detected saccades, including all categories, demonstrates an unimodal pattern with a peak value of approximately 30-40 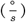 for peak velocity. 50% of detected microsaccades are within the range of 25 to 50 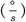 in peak velocity. **F)** Saccade rate amplitude by peak velocity: The distribution of saccade amplitude vs the peak velocity across all categories illustrates a positive linear relationship.

### Microsaccade Detection

Saccade detection commenced with analyzing eye movements for one eye. To enhance the data quality saccades with amplitudes exceeding 2.5 standard deviations from the mean amplitude—for each participant and stimulus category—were considered outliers and excluded from the analysis (Craddock et al., 2017). The selection process for microsaccades then applied specific velocity and amplitude thresholds, distinguishing between slower fixational movements and the rapid eye movements characteristic of microsaccades. Indeed, the velocity thresholds for detecting microsaccades were set using a 6-fold median velocity threshold. Through this approach, we identified potential saccades and isolated microsaccade candidates based on their rapid changes in gaze direction and velocity. Additionally, saccades with an amplitude lower than 1.0° were categorized as microsaccades.

In detecting microsaccades, we further examined amplitude, peak velocity, and direction characteristics. A linear correlation criterion was employed to accurately validate microsaccades, particularly to address overshoot components following corrective fixations post-saccades (Mergenthaler & Engbert, 2010). To maintain accuracy, corrective movements following microsaccades were not considered new microsaccades if they occurred within 30 ms of their preceding one (Møller et al., 2002). Moreover, trials in which subjects did not perform a saccade during the specified period for any stimulus were removed from our analysis to ensure its reliability (Craddock et al., 2017). When all trials are included, our original analysis’s trends and conclusions remain true. This indicates that our approach of excluding non-microsaccade trials did not introduce any significant bias (The analysis is available in Supplementary Information (SI), Section S1).

The saccade detection algorithm is applied to the whole duration of the gaze recordings. We analyze the velocity profile of the gaze data across the entire recording session to ensure robust and accurate detection. In addition, the saccade rate—the number of saccades per second within a trial—was calculated in 15 ms bins. Preprocessing of eye movement data was essential to remove noise and artifacts, enabling accurate microsaccade detection. This preprocessing involved data filtering, blink removal, and baseline drift correction, followed by employing Engbert’s criteria (Engbert & Kliegl, 2003; Mergenthaler & Engbert, 2010) for microsaccade detection during the stimulus presentation period.

### Discriminatory Analysis

We initially applied the detection methodology outlined in Section 2.6 to identify saccades and microsaccades within each trial. Each trial, lasting 900 ms, was analyzed in 15 ms intervals, yielding 60 data points stored in an array per trial. Of these, the first 20 values represent the baseline period, while the subsequent 40 values correspond to the post-stimulus period. This detection process was systematically conducted across all trials for each participant within a session.

#### Statistical Analysis

In our study, we used two statistical methods to analyze the impact of visual stimuli on saccade metrics. First, we employed a one-way repeated measures ANOVA to examine the effects of visual stimuli on microsaccade rate. This method allowed us to account for within-subject variability by treating each subject as their control, thus providing a robust analysis of the influence of different visual stimuli on saccade dynamics. Specifically, we tested the hypothesis that other categories of visual stimuli would result in significant differences in microsaccade rate. The repeated measures post-hoc tests with Bonferroni correction followed ANOVA to identify specific pairs of stimulus categories with significant differences.

Second, we used a bootstrapping approach with ANOVA to compare microsaccade rates across different stimulus categories. Bootstrapping involved repeatedly resampling the data with replacement to estimate the distribution of the sample statistics, thus providing more reliable confidence intervals and significance tests. This approach enabled us to assess whether microsaccade rates significantly differed among the various stimulus categories with greater statistical power and accuracy. For the ANOVA tests, we computed the F-statistics and p-values to quantify the strength and significance of the observed effects. These tests were used to compare microsaccade rates across different stimulus categories. For bootstrapping, we generated confidence intervals for the means and differences in microsaccade rates across categories, ensuring a robust comparison. To elucidate the relationships between visual stimuli and saccade characteristics, such as rate, peak velocity, amplitude, and microsaccade duration, we provided a detailed articulation of our hypotheses and a careful interpretation of the results.

#### Separability Measure

To assess the discriminative capability of microsaccade rates across different visual stimulus categories, we computed a separability measure (*J*). This measure is the ratio of the between-class scatter matrix to the within-class scatter matrix. The separability measure (*J*) was calculated for each category within the designated time window across the entire dataset. We computed *J* values for different time windows to analyze the post-stimulus effect on microsaccade rates across different stimulus categories. For both human and monkey experiments, we reported the *J* values in the 100-500 ms interval.

#### Classification

In the classification task, we utilized a specific 200 ms time window, ranging from -300 ms to 600 ms, with a step size of 5 ms. The results at a given time point *t* represent the window analysis from 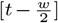 to 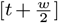 (where *w* is 200 ms in this case). Therefore, the time point *t* refers to the mid-point of the 200 ms window. For example, if a result is reported at time point [*t* = 100] ms, it indicates the analysis of the data within the window [0 ms, 200 ms].

Within this window, we employed an SVM classifier with a radial basis function (RBF) kernel trained on 80% of the dataset (the training set). The classifier’s performance, including recall and accuracy, was evaluated on the remaining 20% of the data (test dataset). For multi-category classification tasks (involving animal, human, man-made, and natural categories), a one-versus-rest strategy was implemented.

### Pupil Size Analysis

In our pupil size analysis, pupil size was recorded simultaneously with eye-tracking data using the same equipment and procedures previously described for monitoring eye movements in human and animal subjects. We performed an ANOVA analysis to assess potential variations in pupil size that could affect microsaccade rates. This analysis compared pupil sizes across different stimulus categories in both the human and monkey datasets. The objective was to identify any significant differences in pupil size that may be associated with or could explain the observed variations in microsaccade rates.

### Neural Data Analysis

In our study, neural data from the inferior temporal (IT) cortex and monkeys’ eye movement data were analyzed. The analysis included data from 157 neurons in Monkey 1 and 109 neurons in Monkey 2, sorted using the ROSS toolbox as outlined in (Toosi et al., 2021).

During our experiments, neural responses to each stimulus were represented as N-dimensional vectors, with each dimension corresponding to the average firing rate of a neuron within a 50 ms window centered on the stimulus presentation. This data was processed using custom Matlab scripts. This involved normalization of neural responses (z-score) and using a sliding window analysis (50 ms window with a 5 ms step) to maintain temporal consistency across trials. The classification of stimulus categories was performed using linear discriminant analysis (LDA) with neural data at the population level. Moreover, the effectiveness of the classifications was compared against saccade-based classifications. We employed a leave-p-out strategy with 30Peak classification timings were examined to explore potential neural feedback effects on microsaccade rates. Additional details on these analytical techniques and their implications are available in (Toosi et al., 2023; Farhang et al., 2021).

## Results

### Experimental paradigm and microsaccade characteristics

To study the impact of stimulus categories on microsaccade rates, we conducted two experiments: the first with two rhesus monkeys and the second with human participants. We utilized the RSVP protocol to gather eye movement data from participants (Fig. 1(A) & Fig. 1(B)), examining microsaccade dynamics across different stimulus categories. The stimulus set contains two superordinate categories, animate and inanimate, and within these, four mid-level categories: animal, human, man-made, and natural scenes. Animals and humans fall under the animate category, while man-made and natural belong to the inanimate category. Moreover, under animate, we considered two mid-level categories: face and body. The face category includes faces from both the animal and human categories, and the body category comprises bodies from the animal and human categories. Throughout our experiments, the baseline period spans from -300 to 0 milliseconds, and the post-stimulus period covers 0 to 600 milliseconds, regarding the stimulus onset time. Our main objective is to explore the relationship between microsaccade patterns and stimulus categories.

First, we analyzed the distribution of saccade amplitudes, focusing on their temporal characteristics and the microsaccade rate, illustrated in Fig. 1(C). It displays the distribution of saccade amplitude overtime during the experiment. The saccade amplitudes mainly ranged from 0.05^*0*^ to 10.2^*0*^ of visual angle. The figure indicates an initial decrease in the saccade rate followed by a subsequent increase in the interval of 200 to 400 ms after stimulus presentation. The saccade rate within the 100–400 ms time window and for amplitudes ranging from 0.1-1.9 degrees exhibit is significantly greater than the baseline period, consequently demonstrating the significant impact of stimulus presentation on the saccade rate (*p*_*value*_ *<* 0.001).

The eye movement data from a monkey participant and a human participant in a sample session are illustrated in Fig. 1(D) and Fig. 1(E), respectively (The gaze coordinates of a human data is also included in SI, Figure S2.). The figures illustrate the participants’ focused gaze movements toward the presented stimulus rather than in other directions.

To validate the detected saccades, it is necessary to assess their distribution across key parameters: time, amplitude, duration, and peak velocity. Fig. 2(A) illustrates the saccade probability and amplitude distribution, demonstrating that 26.4% of detected saccades exhibit amplitudes below 1.0°. The peak of detected saccades during the experiment corresponded to an amplitude of 1.1°. This amplitude threshold determines our targeted movements; those with amplitudes below 1.0° are considered microsaccades. Amplitudes of 69.75% of microsaccade detected in this study ranged between 0.1° and 0.95° of visual angle.

To analyze the saccade distribution over time, we calculate the saccade rate for every 15-millisecond time window across all categories obtained from monkey data, as depicted in Fig. 2(B). The figure presents an unimodal distribution of the saccade rate over time, demonstrating a decreasing trend followed by an increase to its peak after stimulus onset; subsequently, the rate gradually decreases until the end of the trial. On average, the peak of the saccade rate hits 3.35 *±* 0.28 Hz 395 milliseconds after the stimulus onset. Conversely, the minimum saccade rate occurs at 0.47 Hz, observed at 225 milliseconds during the post-stimulus period. The observed increase in the saccade rate following the presentation of stimulus indicates a significant influence of the stimulus presentation on the saccade rate compared with the baseline period (*p*_*value*_ *<* 0.001).

Similarly, Fig. 2(C) illustrates the saccade rate distribution across all categories observed in the human participants. Comparable to the saccade rate distribution observed in monkeys, the temporal pattern of the saccade rate in humans exhibits a similar pattern. The saccade rate peaks at an average of 3.31 *±* 0.32 Hz at 380 milliseconds after the stimulus onset, while the minimum rate is observed at 0.40 Hz at 210 milliseconds after the stimulus presentation.

Visualizing the relationship between microsaccade duration and rate provides insights into the temporal aspects of microsaccadic behavior. Fig. 2(D) illustrates the distribution of microsaccade probability against microsaccade duration. The figure presents an unimodal pattern where higher microsaccade rates correspond with shorter durations (Pearson correlation coefficient, *r* = − 0.70, *p*_*value*_ *<* 0.001). Specifically, the microsaccade rate peaks at 0.28, corresponding with a microsaccade duration of 13 milliseconds. Indeed, the distribution shows an ascending trend leading up to its peak, followed by a decline in value for longer microsaccade durations. Furthermore, the distribution of microsaccade probability concerning their peak velocity is illustrated in Fig. 2(E). The average peak value of the microsaccade rate is 0.11, corresponding to a peak velocity of 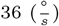.

Previous studies show that the peak velocity of saccades typically increases with an increase in saccadic amplitude. This relationship between peak velocity and saccade amplitude is typically called the “main sequence” (Buonocore et al., 2016). We derive the main sequence by analyzing saccades individually for each participant and then aggregating the data, which is illustrated in Fig. 2(F). The figure demonstrates a statistically significant positive linear correlation between saccade amplitudes and their peak velocities (*R*^2^ = 0.65, *p*_*value*_ *<* 0.001). This relationship suggests that larger amplitudes are associated with higher peak velocities in saccades, as determined by conventional least-square fitting. In the subsequent sections, we will analyze microsaccade rate distributions and statistics across different stimulus categories to examine how various stimulus categories influence microsaccade rates.

### Microsaccade Rate Discrimination Capability

As an initial step to explore the stimulus category impact on the microsaccade rate, we examined the temporal saccade rate distribution across the categories. Fig. 3(A) illustrates the distribution of saccade rates over time for categories (animate and inanimate; face and body; animal and human; man-made and natural). The subfigures presented in Fig. 3(A) indicate a consistent pattern across the categories, characterized by decreased saccade rates followed by an increase in saccade rates reaching their peaks.

**Fig. 3.**
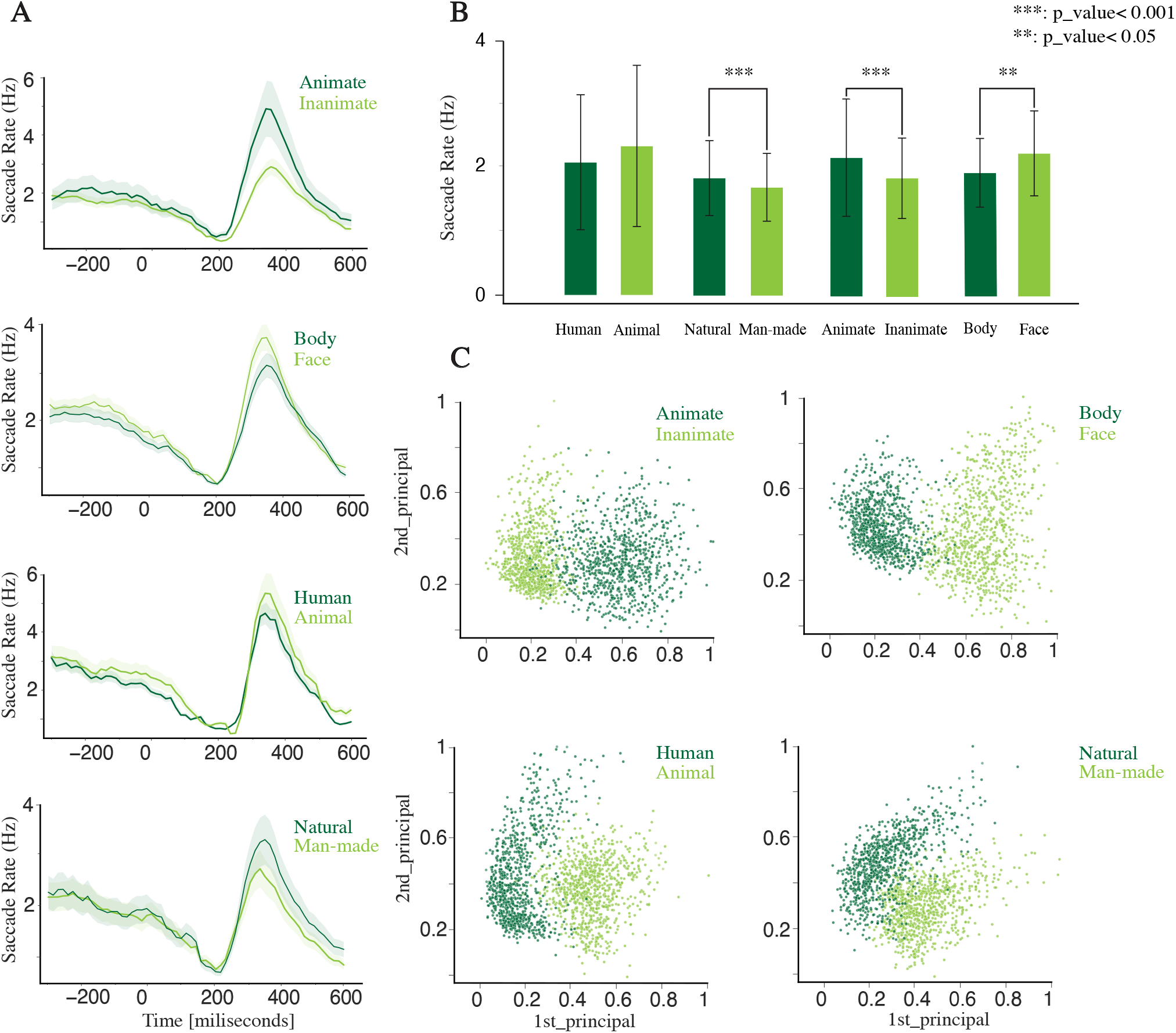
Distinctive Saccade Rate Patterns Among Categories for Monkeys dataset. **A)** Temporal distribution of saccade rate of different categories: The saccade rate distribution per stimulus category reveals an increase in rates in 220 ms post-stimulus presentation, characterized by an unimodal distribution. **B)** Saccade rates statistics across different categories over the entire trial time (900 ms): The mean saccade rates, represented with error bars, exhibit distinct patterns across multiple categories, including animal, human, man-made, natural, animate, inanimate, face, and body stimuli, indicating the distinctness and discernibility of these categories from one another. **C)** Category discrimination through the 2D-PCA representation of microsaccade rates: This figure demonstrates the microsaccade rates’ two-dimensional PCA representation across various stimulus categories. Extracted from their saccadic responses within a time window of 100-600 ms post-stimulus onset. It shows distinct separability among microsaccade rate distributions across different stimulus categories.

In analyzing saccade rate distribution across stimulus categories, we observe distinct variations in the minimum and maximum saccade rates among stimulus categories, particularly during the post-stimulus period. The details of saccade rate statistics for each category are summarized in Table 1.

**Table 1.**
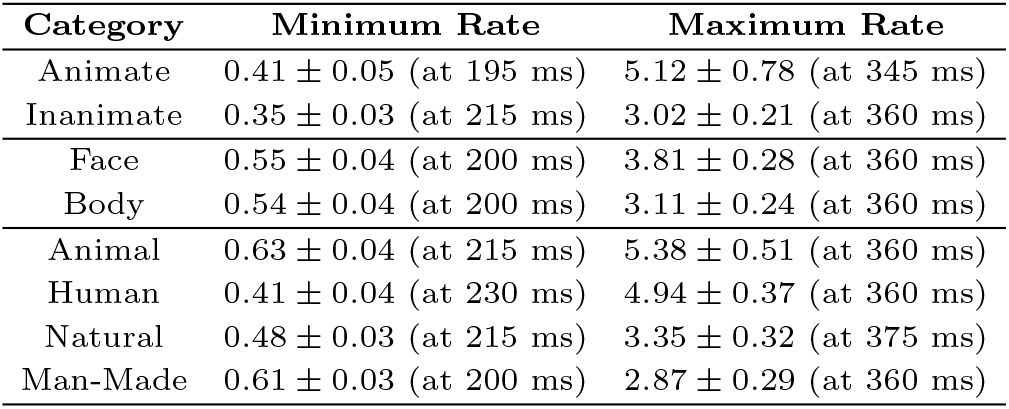
Saccade Rate Statistics separated by Category in Monkey.

| Category | Minimum Rate | Maximum Rate |
| --- | --- | --- |
| Animate | $0.41 \pm 0.05$ (at 195 ms) | $5.12 \pm 0.78$ (at 345 ms) |
| Inanimate | $0.35 \pm 0.03$ (at 215 ms) | $3.02 \pm 0.21$ (at 360 ms) |
| Face | $0.55 \pm 0.04$ (at 200 ms) | $3.81 \pm 0.28$ (at 360 ms) |
| Body | $0.54 \pm 0.04$ (at 200 ms) | $3.11 \pm 0.24$ (at 360 ms) |
| Animal | $0.63 \pm 0.04$ (at 215 ms) | $5.38 \pm 0.51$ (at 360 ms) |
| Human | $0.41 \pm 0.04$ (at 230 ms) | $4.94 \pm 0.37$ (at 360 ms) |
| Natural | $0.48 \pm 0.03$ (at 215 ms) | $3.35 \pm 0.32$ (at 375 ms) |
| Man-Made | $0.61 \pm 0.03$ (at 200 ms) | $2.87 \pm 0.29$ (at 360 ms) |

**Table 2.** Classification Results Summary in Monkey.

| Comparison | Baseline Accuracy | Peak Accuracy (Time) | End Accuracy | Recall Values |
| --- | --- | --- | --- | --- |
| Animate vs. Inanimate | 51.67% ± 0.24 | 79.52% ± 0.36 (315ms) | 52.72% ± 0.21 | 79.32%, 78.54% |
| Face vs. Body | 50.92% ± 0.24 | 75.85% ± 0.32 (315ms) | 53.47% ± 0.18 | 77.28%, 76.24% |
| Human vs. Animal | 50.77% ± 0.29 | 77.68% ± 0.45 (300ms) | 52.72% ± 0.21 | 79.18%, 78.47% |
| Man-made vs. Natural | 50.26% ± 0.33 | 76.07% ± 0.38 (300ms) | 51.95% ± 0.27 | 77.49%, 76.65% |
| All Categories | 26.11% ± 0.39 | 72.75% ± 0.49 (315ms) | 52.65% ± 0.22 | Animal: 77.38%, Human: 75.52%,<br>Man-made: 76.44%, Natural: 77.39% |

Analysis of the saccade rate statistics from Table 1 and Fig. 3(B) reveals distinct patterns in saccadic response dynamics across different stimulus categories. The statistics highlight that the animal category exhibits the highest mean saccade rate among the stimulus categories, followed by human, natural, and man-made stimuli. At the superordinate level, the animate category demonstrates a higher average saccade rate than the inanimate category. Furthermore, the face category has a higher mean saccade rate than the body category.

Significant distinctions in microsaccade rate (MSR) are also observed across categories, between the animate and inanimate (*MSR*_*animate*_ = 1.04 *±* 0.13, *MSR*_*Inanimate*_ = 0.81 *±* 0.09, one-way ANOVA *p*_*value*_ *<* 0.001, *F*_*statistic*_ = 63.48) as well as face and body (*MSR*_*face*_ = 0.9 *±* 0.15, *MSR*_*body*_ = 0.65 *±* 0.11, one-way ANOVA, *p*_*value*_ *<* 0.05, *F*_*statistic*_ = 54.18). Moreover, within the mid-level categories, the stimulus categories exhibit significant differences from each other (*MSR*_*animal*_ = 1.2 *±* 0.11, *MSR*_*human*_ = 1.01 *±* 0.09, *MSR*_*man-made*_ = 0.72 *±* 0.11, *MSR*_*natural*_ = 0.86 *±* 0.13, one-way ANOVA, *p*_*value*_ *<* 0.001, *F*_*statistic*_ = 93.33). The bar plot of microsaccade rate statistics for the monkey dataset is also included in SI, Figure S5 (A).

Given the statistically significant difference in microsaccade rates across categories, we aimed to further investigate category discrimination based on these rates within a time window of 50-600 milliseconds post-stimulus onset. To quantify the distinctiveness of stimulus categories, we apply Principal Component Analysis (PCA) and separability measure (J), focusing on their microsaccadic rate distributions in 100-600 milliseconds post-stimulus. Fig. 3(C) illustrates a two-dimensional PCA visualization (using the first two principal components) that contrasts pairs of categories: animate vs. inanimate, face vs. body, animal vs. human, and man-made vs. natural categories. The resulting scatter plots exhibit distinct clusters for each category pair, visually highlighting their differentiation. Furthermore, Section S4 provides the analysis of a few of the first principal components, corresponding explained variances, and PCA on the entire dataset, including all stimulus categories.

Analysis of calculating the separability measure values (J) across different categories indicates significant distinction, particularly within the 100 to 500 milliseconds post-stimulus window, exhibiting this period as the highest degree of separability (27.88 for animate versus inanimate, 24.32 for face versus body, and 20.2 across all sub-categories). Following this, an SVM will be employed to examine the hypothesis concerning the selectivity of microsaccade rates by stimulus category, with the results detailed in the following section.

### Classification of Stimulus Categories Based on Microsaccade Rates

To evaluate the effectiveness of using microsaccade rates for the classification of stimulus categories, SVM is employed across discrete 200-millisecond intervals within the saccade rate distribution to classify the categories, starting from the baseline period (−300-0 milliseconds) to the end of the post-stimulus period (0-600 milliseconds). Fig. 4 displays the classification results, illustrating accuracy and recall metrics for various categories. The accuracy increases during the post-stimulus period, reaching a peak at around 300-320 milliseconds after stimulus onset. The high accuracy values are consistently observed within 200 to 400 milliseconds following the stimulus onset. Hence, the recall values, alongside error bars, are reported for this time window (200-400 milliseconds post-stimulus period). The classification results for each comparison (animate versus inanimate, face versus body, human versus animal, man-made versus natural, and across all stimulus categories) are illustrated in Fig. 4(A-E).

The classification results, detailed in Table 2, demonstrate significant discrimination capability of microsaccade rate and its variations in classification across different stimulus categories within the specified post-stimulus time window of 100-600 milliseconds. These results indicate that stimulus categories at all hierarchical levels (superordinate and mid-level) can be distinguished with an accuracy exceeding 70.0%. This result shows that the distinctions between animate and inanimate, face and body, as well as among stimulus categories (animal, human, man-made, and natural), are achievable through analysis of their post-stimulus microsaccade rate distributions.

To ensure our classification model’s robustness and reliability, we analyzed a trial-shuffled dataset, where any inherent relationships between the microsaccade data and the stimulus categories are disturbed. As anticipated, the performance metrics, including accuracy and recall, dropped significantly in the trial-shuffled dataset compared to the original dataset. This substantial decrease confirms that the classifier’s high performance with the true labels is not due to random chance. Detailed results of this analysis can be found in the Section S5.

### Microsaccade Rate Analysis and Category

#### Classification in Humans

To explore the influence of stimulus presentation on saccade rates within the human data, we employ an analytical approach similar to that used for the monkey data. Fig. 5(A) represents the detected saccade distribution across different categories (animate vs. inanimate, face vs. body, animal vs. human, and natural vs. man-made), mirroring patterns observed in the monkey data. The distribution of saccade rates across the category reflects an unimodal pattern over time. In the analysis of saccade rate distributions across the stimulus categories, distinct variations are observed in the minimum and maximum saccade rates during different post-stimulus periods, detailed in Table 3.

Consistent with previous results (Fig. 3, Table 1), the analysis of saccade rate, as shown in Table 3 and Fig. 5(B), exhibits a distinct saccadic response pattern across different stimulus categories. The categories achieving the highest maximum saccade rates include the animate category at the superordinate level and the face and the animal categories at the mid-level, surpassing all others followed by human, natural, and man-made stimuli.

**Table 3.**
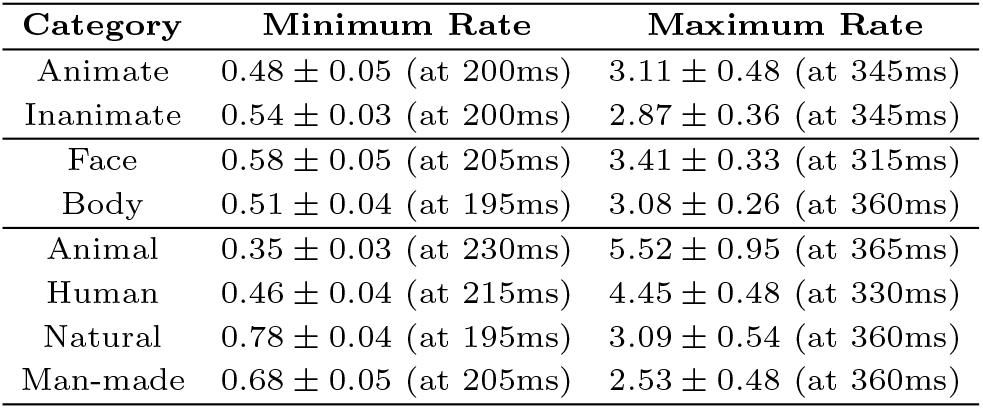
Saccade Rate Statistics Separated by Category in Human.

| Category | Minimum Rate | Maximum Rate |
| --- | --- | --- |
| Animate | $0.48 \pm 0.05$ (at 200ms) | $3.11 \pm 0.48$ (at 345ms) |
| Inanimate | $0.54 \pm 0.03$ (at 200ms) | $2.87 \pm 0.36$ (at 345ms) |
| Face | $0.58 \pm 0.05$ (at 205ms) | $3.41 \pm 0.33$ (at 315ms) |
| Body | $0.51 \pm 0.04$ (at 195ms) | $3.08 \pm 0.26$ (at 360ms) |
| Animal | $0.35 \pm 0.03$ (at 230ms) | $5.52 \pm 0.95$ (at 365ms) |
| Human | $0.46 \pm 0.04$ (at 215ms) | $4.45 \pm 0.48$ (at 330ms) |
| Natural | $0.78 \pm 0.04$ (at 195ms) | $3.09 \pm 0.54$ (at 360ms) |
| Man-made | $0.68 \pm 0.05$ (at 205ms) | $2.53 \pm 0.48$ (at 360ms) |

Moreover, significant distinctions in MSR are detected across categories. Specifically, a significant difference is observed between the animate and inanimate (*MSR*_*animate*_ = 1.06 *±* 0.08, *MSR*_*Inanimate*_ = 0.82 *±* 0.06; one-way ANOVA, *p <* 0.001, *F*_*statistic*_ = 82.62), and between the face and body (*MSR*_*face*_ = 0.95 *±* 0.11, *MSR*_*body*_ = 0.69 *±* 0.08; one-way ANOVA, *p <* 0.001, *F*_*statistic*_ = 60.08). Additionally, significant differences are found across all stimulus categories (*MSR*_*animal*_ = 1.3 *±* 0.12, *MSR*_*human*_ = 1.08 *±* 0.08, *MSR*_*man-made*_ = 0.79 *±*0.07, *MSR*_*natural*_ = 0.87 *±*0.06; one-way ANOVA, *p <* 0.001, *F*_*statistic*_ = 102.12), exhibiting the distinct microsaccadic dynamics characteristic of each category. The bar plot of microsaccade rate statistics for the human dataset is also included in SI, Figure S5 (B).

Furthermore, the analysis of category separability, using the separability measure, J, within the 100-500 milliseconds post-stimulus time window, reveals significant distinctiveness across categories. Specifically, the J values for comparisons between animate versus inanimate, face versus body, and across all categories are 22.41, 21.25, and 18.33, respectively, indicating a high degree of discriminability.

Similar to the previous experiment with monkeys (Fig. 4), Fig. 6 illustrates the classification accuracy and mean recall values in human experiment. High accuracies are consistently observed in the 200 to 400 milliseconds post-stimulus onset. In addition, detailed classification results for each comparison are illustrated in Fig. 6(A) through (E), corresponding to animate/inanimate, face/body, human/animal, natural/man-made, and across all stimulus categories, respectively. The classification results specifications are provided in Table 4.

**Table 4.**
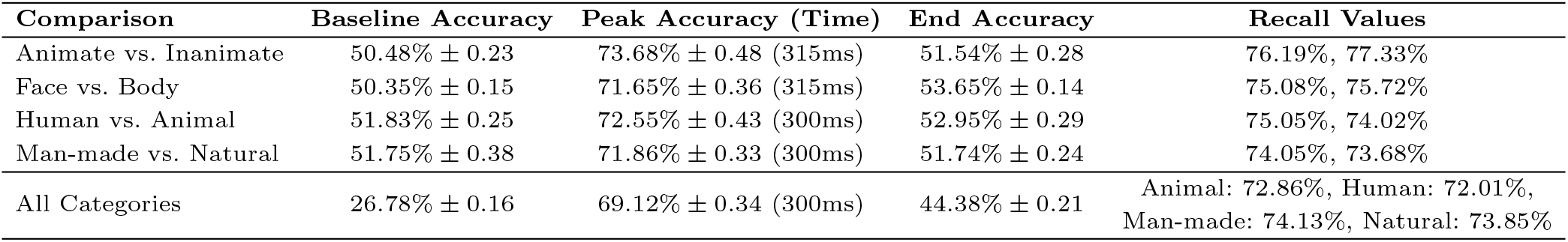
Classification Results Summary in Human.

**Fig. 4.**
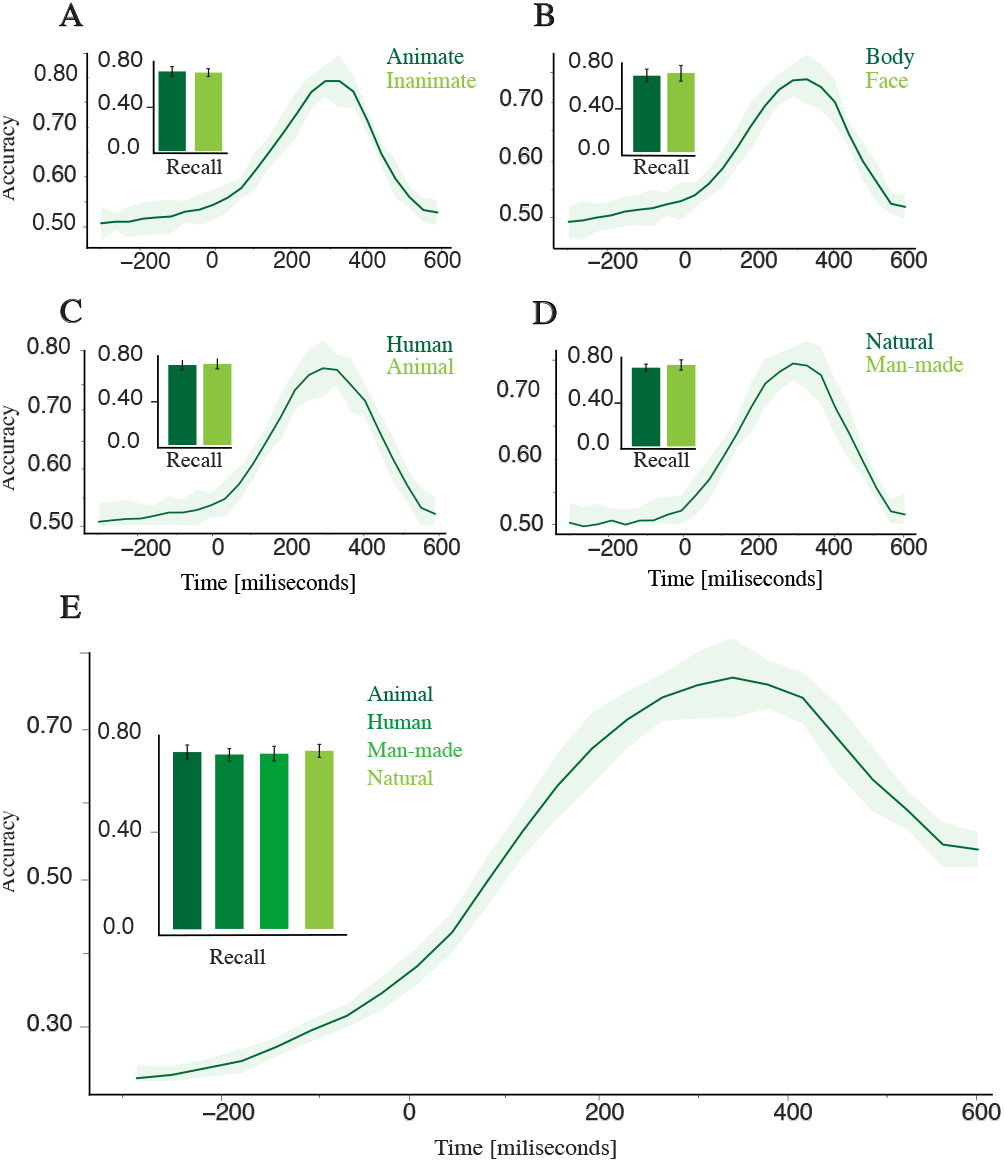
Time course of the performance of classifying categories with microsaccade rate employing an SVM classifier for monkeys dataset. The mean and standard deviation of classification accuracy are illustrated for all cases throughout each trial. The accuracy distributions are unimodal, with increasing accuracy after stimulus onset and peaking around 200-400 ms after stimulus presentation. The bar plots depicted in each figure present the mean and standard deviation of the recall values within the time interval of 200 - 400 ms following the stimulus onset. **A)** Animate vs. Inanimate: The accuracy of classifying between animate and inanimate categories has the highest accuracy, peaking at approximately 84%. **B)** Face vs. Body: The classification accuracy for classifying between face and body categories has peak accuracy reaching approximately 75%. **C)** Human vs. Animal: The classification accuracy between animal and human categories reaches its maximum accuracy at around 79%. **D)** Man-made vs. Natural: The classification accuracy between man-made and natural categories has the highest accuracy reaching approximately 76%. **E)** All categories: The classification accuracy for classifying between all categories reaches its highest point at approximately 74%.

**Fig. 5.**
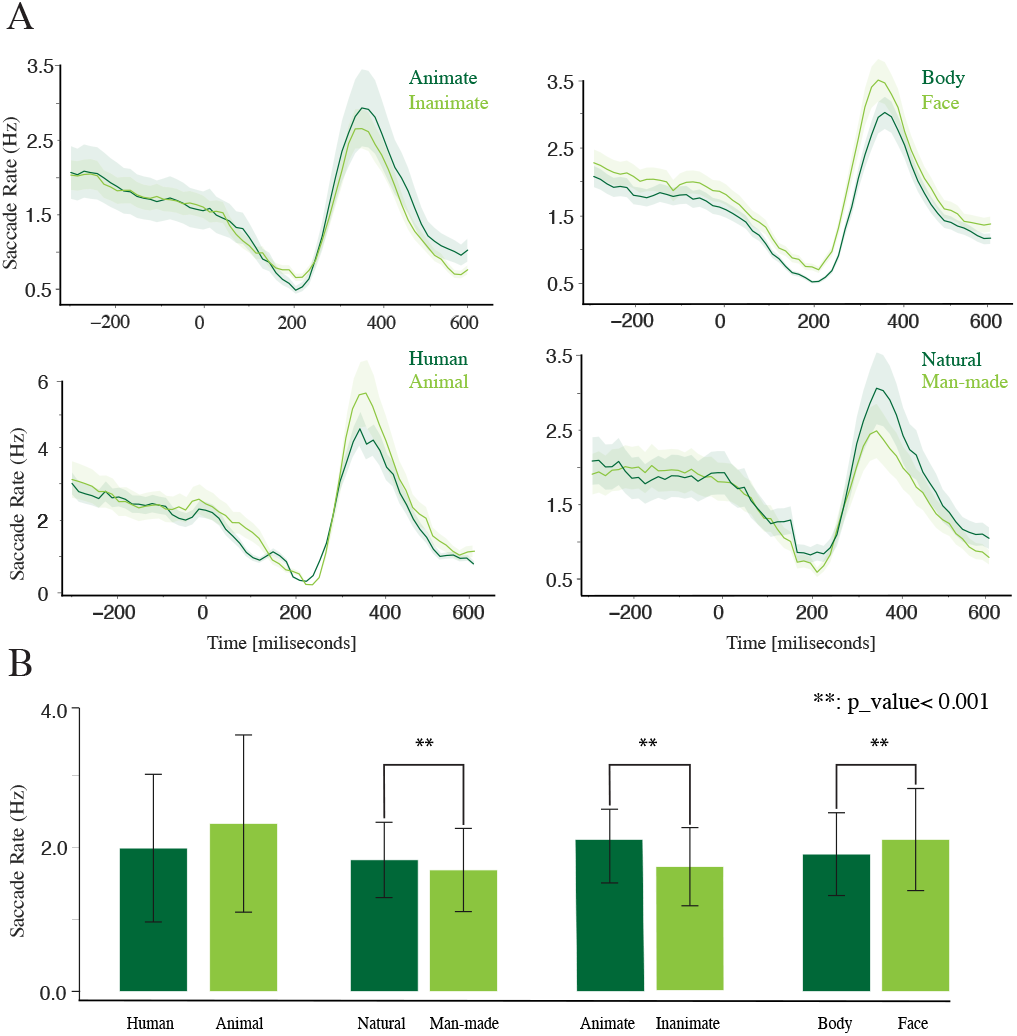
Distinctive microsaccade rate patterns. **A)** Saccade rate temporal distribution of different categories for the human dataset: The distribution of saccade rates for each stimulus category observed in the human data reveals a rise in saccade rates approximately 200 ms after stimulus presentation, characterized by an unimodal distribution primarily influenced by the stimulus presentation. Each bin represents a 15 ms interval. **B)** Saccade rates statistics across different categories over the entire trial time (900 ms): The mean saccade rates for various categories are represented with error bars indicating standard deviation, including all stimuli collected from human data, and exhibit variability across multiple categories.

**Fig. 6.**
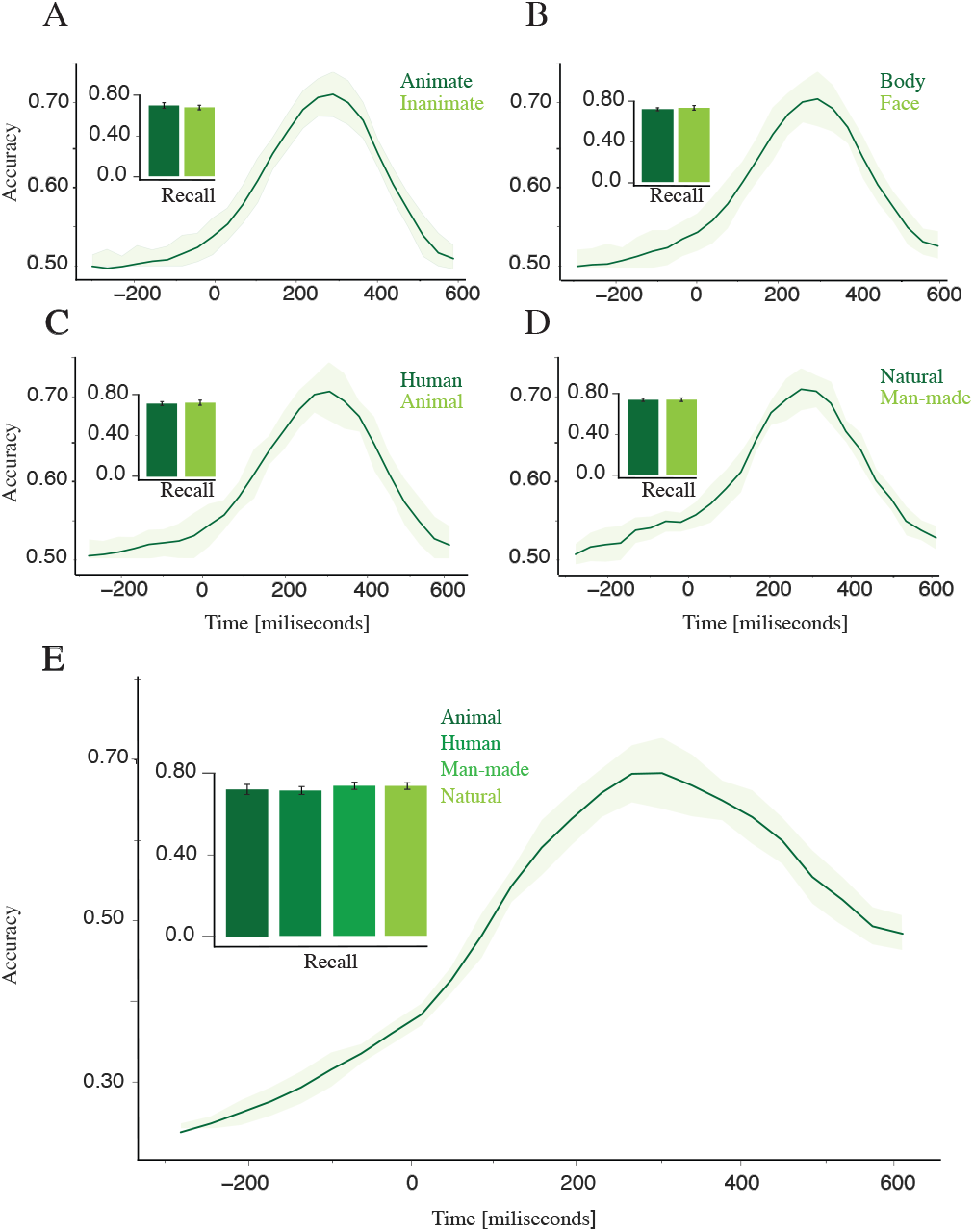
Classification of the stimulus categories in the human dataset. The mean and standard deviation of classification accuracy and recall value for all cases throughout each trial using a linear classifier. **A)** Animate vs. Inanimate: Classifying between animate and inanimate categories has the highest accuracy, peaking at approximately 73%. **B)** Face vs. Body: The classification accuracy for classifying face and body categories has peak accuracy reaching approximately 71%. **C)** Human vs. Animal: The accuracy of classifying between animal and human categories reaches its maximum accuracy at around 71%. **D)** Man-made vs. Natural: Classifying between man-made and natural categories has the highest accuracy, reaching approximately 72%. **E)** The classification accuracy between all categories reached its highest point at approximately 69%.

Like the monkey experiment, the classification results reveal significant variability in classification metrics across distinct stimulus category comparisons within the particular post-stimulus window of 100-600 milliseconds. These observations indicate that stimulus categories across all hierarchical levels are discernible by microsaccade rate, with classification metrics surpassing 70.0%. Notably, the analysis of post-stimulus microsaccade rate distributions enables the clear differentiation between animate and inanimate, face and body, and distinctions among all stimulus categories.

In pairwise decoding of both experiments, the end accuracy approaches the baseline level, indicating that the transient microsaccade responses specific to each pair diminish over time. Conversely, in decoding across all categories, the end accuracy remains above chance, as the aggregated microsaccade patterns from multiple categories retain discriminative features longer due to the richness of the combined dataset in multi-category decoding, which helps maintain accuracy values higher than the chance level. Detailed analyses and figures supporting these observations are provided in Section S2.

In the following section (Section 4), we conduct additional analyses incorporating pupil size and neural data to explore further the underlying factors influencing microsaccade rates in response to different visual stimuli.

## Brain State and Neural Feedback in Microsaccades

This section aims to determine the underlying factors influencing microsaccade rates in response to different visual stimulus categories. Specifically, we aim to explore whether the observed differences in microsaccade rates across stimulus categories are related to the brain state, such as pupil size, or are influenced by neural feedback mechanisms following stimulus discrimination in the IT cortex.

Understanding the origin of microsaccade rate differences is crucial for interpreting their role in visual processing and object categorization. If these differences are related to brain states, such as changes in pupil size, it would suggest a link to overall cognitive or attentional states. Conversely, if neural feedback mechanisms influence the differences, it would imply a more direct role of specific neural circuits in modulating eye movements based on visual input. To address these questions, we conducted two separate analyses:

1. **Pupil Size Analysis:** We compared the pupil sizes across different stimulus categories for both human and monkey datasets to determine if there were significant differences that could account for the observed microsaccade rate variations.
2. **Neural Data Analysis:** We compared the accuracy of stimulus category classification using saccade data and neural data from the IT cortex. We focused on the timing of peak classification accuracy to determine if neural feedback mechanisms might influence microsaccade rates.

Pupil Size Analysis

Fig. 7 (A, B) compares pupil sizes across different stimulus categories for both monkey (Fig. 7(A)) and human (Fig. 7(B)) datasets. The results indicate that pupil size is not significantly different across the categories, suggesting that the differences in microsaccade rates are unrelated to brain state as measured by pupil size.

**Fig. 7.**
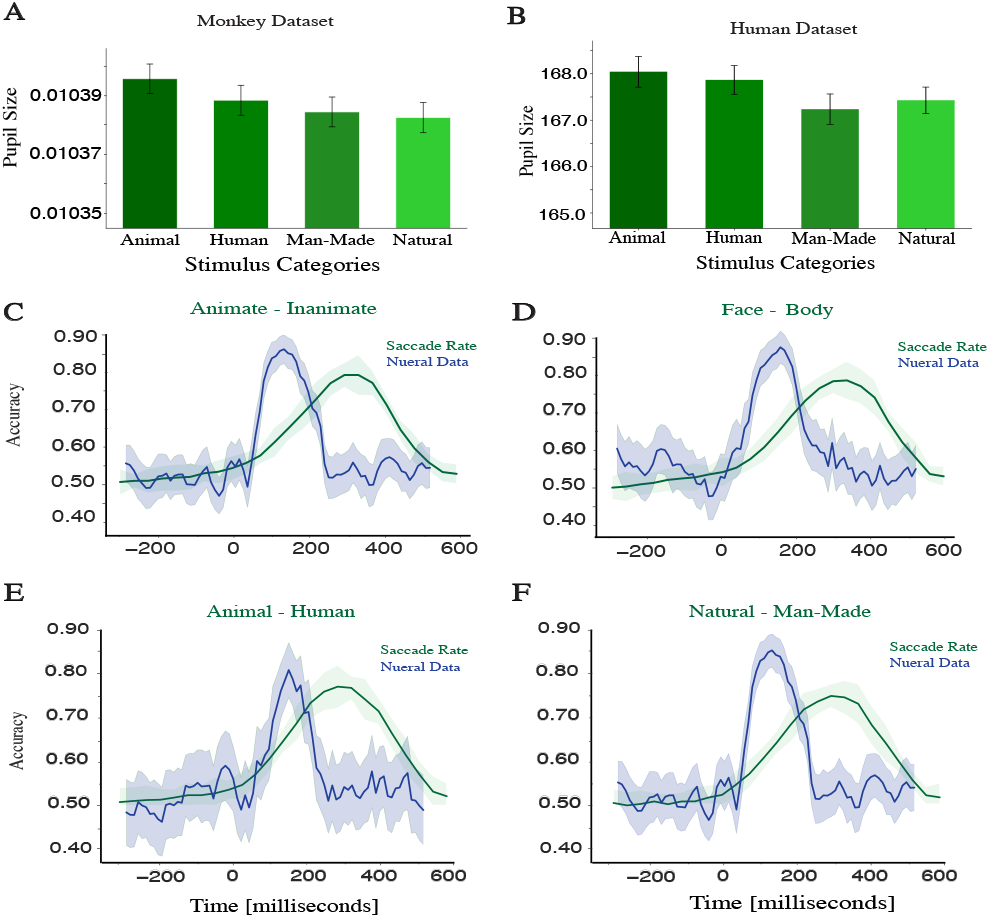
Pupil size comparison and Classification accuracy over time with neural data & saccade rate. **A)** Pupil size comparison across different stimulus categories for the monkey dataset. **B)** Pupil size comparison across different stimulus categories for the human dataset. **C)** Classification accuracy over time with neural data & saccade rate for Animate vs. Inanimate stimulus categories. **D)** Classification accuracy over time with neural data & saccade rate for Face vs. Body stimulus categories. **E)** Classification accuracy over time with neural data & saccade rate for Animal vs. Human stimulus categories. **F)** Classification accuracy over time with neural data & saccade rate for Natural vs. Man-Made stimulus categories.

For the monkey dataset, the mean values for pupil sizes are as follows: Animal (0.010395 *±* 0.00001), Human (0.010388 *±* 0.00001), Man-Made (0.010384 *±* 0.00001), and Natural (0.010382 *±* 0.00001). The ANOVA results showed no significant difference (*F*_*statistic*_ = 1.427,*p*_*value*_ = 0.232). For the human dataset, the mean values for pupil sizes are as follows: Animal (168.08 *±* 0.75), Human (167.82 *±* 0.71), Man-Made (167.25*±*0.73), and Natural (167.42*±*0.68). The ANOVA results showed no significant difference (*F*_*statistic*_ = 1.564, *p*_*value*_ = 0.195). The analysis of pupil size across different stimulus categories indicates that brain state, as measured by pupil size, does not account for the observed differences in microsaccade rates.

### Neural Data Analysis

In our experiments, we analyzed the monkeys’ eye movement data from a larger dataset, including neural data recorded from the inferior temporal cortex. For further details on neural data recording, please refer to the works of Toosi et al. (2023) and Farhang et al. (2021). Fig. 7(C-F) presents the accuracy of classifying stimulus categories using saccade data and neural data. These results indicate that neural data can effectively discriminate between different stimulus categories (Animate vs. Inanimate: 0.855*±*0.039 at 115 ms, Face vs. Body: 0.903*±*0.052 at 135 ms, Animal vs. Human: 0.806 *±* 0.065 at 135 ms, Man-Made vs. Natural: 0.855 *±* 0.039 at 115 ms).

When comparing the peak accuracy times of classification with neural data to saccade data (Table 2), the peak accuracy value for discriminating stimulus categories with neural data occurs much earlier in the IT cortex than in the saccade data. This result suggests that after the initial discrimination of stimulus categories in the IT cortex, a feedback mechanism influences eye movements, leading to changes in microsaccade rates.

## Discussion

This study explored how different stimulus categories affect saccade rates during a passive viewing task. We found that the category of stimulus significantly influences the microsaccade rates. The distribution of saccade rates showed an initial decrease followed by an increase immediately after the stimulus presentation. We observed that animal stimulus, representing a mid-level abstraction, elicited the highest average microsaccade rates, whereas man-made stimulus resulted in the lowest. At a superordinate level, the animate category (comprising animals and humans) yielded higher microsaccade rates than the inanimate category (including natural and man-made categories). Within the animate category, images depicting faces led to a greater increase in microsaccade rates than those showing bodies, highlighting the impact of facial features on visual attention. Our study demonstrated unique patterns of microsaccade rate distribution following stimulus presentation specific to different categories of stimuli. This suggests the potential for classifying stimuli based on these patterns. Using SVM to classify stimulus categories based on microsaccade rate distributions, we achieved accuracy rates of 70 to 85% in specific trial scenarios within a 200 to 400 ms window after stimulus onset. These results underscore the role of microsaccades as a reliable indicator of object content, reinforcing their significance in object categorization. Comparatively, the difference in saccade rates across categories was less pronounced in the human dataset than in the monkey dataset, leading to a slight decrease in classification performance metrics by 3-6%.

Our results highlighted the influence of stimulus presentation on saccade amplitude, echoing previous research (Kauffmann et al., 2019; Craddock et al., 2017) that demonstrated the modulation of saccade amplitude by the stimulus. Specifically, we observed a shift in the amplitude distribution during the post-stimulus period, with the majority of amplitudes being less than 1.0° (Fig. 1(D)). This shift suggests an increase in microsaccade rates following stimulus presentation, indicating a change in saccade amplitude within this time interval. The observed amplitude distributions affirm the significant impact of stimulus presentation in modulating saccade rate and amplitude. Additionally, the unimodal distribution of saccade probability by amplitude, peaking at 1.1° and declining for higher amplitudes (Fig. 2(A)), aligns with the findings of Pomplun et al. (2001).

Our study elucidated the dynamic relationship between microsaccade peak velocities and stimulus presentation, where peak velocities ranging from 24.2 to 94.72 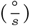, which is 25. in line with the work of Buonocore et al. (2017). Our analysis of the relationship between peak velocity, amplitude, and microsaccade rate revealed that peak microsaccade rates 26. occur between 20 to 40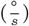. These rates are associated with smaller amplitudes in comparison to microsaccades with higher peak velocities, as depicted in (Fig. 2(E) and Fig. 2(F)). This observation is consistent with the results of Valsecchi et al. (2007), which describe the distribution of microsaccade probability by peak velocity as unimodal. Additionally, our results identified a significant positive linear correlation between microsaccade amplitude and peak velocity (*p*_*value*_ *<* 0.001), resonating the results of Møller et al. (2002) and Krejtz et al. (2018). This correlation is visually supported by Fig. 2(F) and aligns with the works of (Møller et al., 2002; Krejtz et al., 2018; Buonocore et al., 2016).

The impact of stimulus categories on saccade rates is clear through the consistent patterns observed across different stimulus categories. Analysis of saccade rate distributions exhibited an unimodal distribution post-stimulus onset, consistent with previous research findings of Kauffmann et al. (2019), displaying an increase in saccade rate in the post-stimulus period. We observed a distinct decline in microsaccade rate during the post-stimulus period, followed by an increase, with rates peaking between 200-400 milliseconds. This pattern mirrors observations reported in prior research (Craddock et al., 2017; Rosenzweig & Bonneh, 2019; Kadosh & Bonneh, 2022b). Rosenzweig & Bonneh (2019) found that familiar faces exhibited a significantly lower peak than unfamiliar faces. Our findings also highlighted significant differences across stimulus categories (animals, humans, man-made objects, and natural scenes) and at the superordinate level (animate versus inanimate), including distinctions between faces and bodies. These results illustrate the distinct effects of stimulus categories on microsaccade rates, emphasizing the role of microsaccadic activity in visual processing. Complementing this, amplitude analysis conducted by Kauffmann et al. (2019) showed that saccades were larger for face targets than for vehicles, further underlining the influence of stimulus type on saccadic responses. Consistent with this, our analysis revealed that the animal category elicited the highest average microsaccade rate, followed by humans, natural scenes, and man-made objects. Additionally, images of faces induced higher microsaccade rates than those of bodies, as illustrated in Fig. 3 and Fig. 5.

The higher microsaccade rates observed in response to animal and human categories may signify more dynamic and intense visual processing or attentional mechanisms directed toward these stimulus categories, as indicated by Rosenzweig & Bonneh (2019); Kauffmann et al. (2019). Faces are particularly salient visual stimuli, attracting significant attention and interest due to their social significance. Such increased saccade rates in these categories could be indicated by increasing cognitive and visual processing (Kadosh & Bonneh, 2022a; Salvia et al., 2020; Devue et al., 2012), potentially signifying enhanced recognition, emotional processing, or cognitive evaluation in response to these stimuli. Conversely, lower microsaccade rates in other categories, including inanimate objects (man-made and natural) and body images, may reflect differences in cognitive processing, visual interest, or attentional engagement across these categories. This underscores the potential influence of stimulus content in modulating saccadic responses and highlights the complex relationship between visual stimuli and saccadic movements. Our results indicate distinct microsaccade rate patterns across different stimulus categories, revealing differential temporal dynamics in saccadic behavior related to the stimulus categories presented in our experiment.

A significant challenge in interpreting the modulation of microsaccade rates is understanding the underlying factors that drive these changes. Specifically, whether these modulations are related to psychophysical detection, perceptual discrimination, stimulus saliency, or neural feedback mechanisms remains unclear. Our analysis of pupil size revealed no significant differences across different stimulus categories, suggesting that changes in brain state, as indicated by pupil dynamics, are not the primary drivers of the observed microsaccade rate modulation (Fig. 7(A, B)). Furthermore, neural data analyses indicated that the peak accuracy for classifying stimulus categories in the IT cortex occurs earlier than the peak microsaccade rate (Fig. 7(C-F)). These results imply a potential feedback mechanism from the IT cortex influencing eye movements after stimulus discrimination rather than directly correlating with immediate attentional processes. Our results demonstrate that a combination of perceptual and neural processes more likely influences the observed differences in microsaccade rates across stimulus categories. These results suggest that microsaccade rates can serve as a discriminative marker for visual stimulus classification. However, their modulation is potentially linked to neural feedback mechanisms post-stimulus discrimination. Further experiments and studies are necessary to explore the potential relationship between the neural mechanisms in the IT cortex and the microsaccade rate.

To the best of our knowledge, this study is the first to analyze the separability of microsaccade rates across different stimulus categories. Our results demonstrate distinct separability between categories, enabling the differentiation of animal, human, man-made, natural, animate, inanimate, face, and body stimuli based on their microsaccade rate distributions during the post-stimulus period. We utilized an SVM to analyze microsaccade rate data from individual trials, investigating the capability of microsaccades for object decoding. This approach aligns with the application of SVMs to eye-tracking and EEG data to classify the cognitive workload against a no-task control condition (Borys et al., 2017). However, our study focuses exclusively on eye-tracking data, particularly microsaccade rate, to classify stimulus categories.

Moreover, most existing research focuses on exploring the relationship between microsaccades and the emergence of peaks in gamma-band oscillations, typically within the 30 to 100 Hz range, as detected through scalp-recorded EEG. This gamma-band activity is believed to signal the activation of object representations and the synthesis of neural activity from diverse neuron populations, each encoding distinct aspects of the object into a unified percept (Tallon-Baudry & Bertrand, 1999). Consequently, many observed variations in induced gamma-band activity, such as those resulting from changes in object orientation (Martinovic et al., 2007), are likely due to alterations in saccade rate during the critical window of 200–400 ms post-stimulus. This aligns with our results that classification performance metrics for stimulus categories peak within the 200-400 ms period following stimulus presentation.

Our classification analysis (Fig. 4 and Fig. 6) demonstrates the feasibility of differentiating stimulus categories based on microsaccade rates. The significant performance metrics we obtained across various categories highlight the capability to identify distinct stimulus categories through analysis of microsaccade rate patterns. This finding suggests microsaccade rates as potential discriminative markers for visual stimulus classification, marking a step forward in our understanding of microsaccades’ utility as an analytical tool for object categorization. This opens a way for future research in areas ranging from neuroscience to computer vision, with broad implications for cognitive science and artificial intelligence. Our study reveals how visual stimuli content significantly impacts microsaccade rates, providing insights into the dynamic saccadic responses to various stimulus categories. The differences in microsaccade rates across categories highlight the influence of stimulus properties on microsaccadic activities. Furthermore, our research highlights the utility of classification algorithms in discerning stimulus categories through distinct microsaccade rate patterns. Therefore, analyzing saccade rates during the post-stimulus period could provide valuable perspectives on object recognition processes and the contribution of eye movements in processing spatial information.

## Data Availability

The codes and notebooks utilized in this study are deposited on GitHub at https://github.com/sa-nouri/ms-selectivity-feature/tree/main. Furthermore, a few samples of the data examined in this study can be accessed at ms-selectivity-feature-data. All the experimental datasets produced in this study will be accessible upon reasonable request from the corresponding author.

