## Supplementary Information for "Microsaccade Selectivity as Discriminative Feature for Object Decoding"

#### 1. EXCLUDING NON-SACCADES TRIALS

Our current approach removes trials where subjects did not perform a saccade during the specified period for any stimulus. This decision was made to ensure the reliability of our analysis, as trials without microsaccades may not contribute to the understanding of stimulus-driven microsaccade dynamics. This approach is supported by [1], which suggests that excluding such trials can enhance the reliability of microsaccade analysis.

To address concerns about this approach, we conducted additional analyses that included the trials without microsaccades. The results were compared with our original results to ensure the observed trends remained consistent. Microsaccade rate (MSR) was defined as the frequency of microsaccades within a given time window. The overall microsaccade rates were slightly reduced due to including non-microsaccade trials, but the relative differences between stimulus categories persisted. We used the 0-600 milliseconds post-stimulus time window for statistical comparisons about removing the trials without the microsaccade effect. Specifically, the statistically significant differences across the microsaccade rates of different stimulus categories were maintained, as detailed in Table S1.

| **Comparison** | **Statistical Test** | ***p*-value** | ***F*-statistic** |
|---|---|---|---|
| $MSR_{Animate}$ vs. $MSR_{Inanimate}$ | One-way ANOVA | $< 0.001$ | 59.37 |
| $MSR_{Face}$ vs. $MSR_{Body}$ | One-way ANOVA | $< 0.05$ | 50.27 |
| $MSR_{Animal}$ vs. $MSR_{Human}$ vs. $MSR_{Man-Made}$ vs. $MSR_{Natural}$ | One-way ANOVA | $< 0.001$ | 89.84 |

**Table S1.** Summary of ANOVA test results for microsaccade rate comparisons.

We also conducted an ANOVA test to compare the microsaccade rates with and without including non-microsaccade trials, which showed no significant differences between the two sets ($F = 1.05$, $p = 0.31$).

The decoding accuracy using microsaccade rates remained significantly above chance levels within and across superordinate stimulus categories. The patterns observed were similar to those in the original analysis. An ANOVA test was conducted to compare the classification accuracy and predicted labels (stimulus categories) with and without removing the non-microsaccade trials. This comparison was performed under two different scenarios: (1) on the predicted labels (stimulus categories) and (2) on the accuracy values. The results indicated no significant difference in both scenarios. The ANOVA test yielded $F_{statistic} = 1.05$, $p_{value} = 0.31$ for the predicted labels. The ANOVA test yielded $F_{statistic} = 1.20 and p_{value} = 0.27$ for the accuracy values.

These results confirm that our approach of excluding non-microsaccade trials did not introduce any significant bias. The key trends and conclusions of our original analysis remain valid, indicating the robustness of our results.

#### 2. PAIRWISE AND MULTI-CATEGORY DECODING ACCURACY

In the pairwise decoding analysis, the end accuracy returns close to the baseline accuracy. This trend can be attributed to the transient nature of microsaccade responses in each pair of comparisons. Initially, distinct microsaccade patterns post-stimulus facilitate higher decoding accuracy, but as time progresses, these patterns become less pronounced, causing the accuracy to approach the baseline. However, as illustrated in

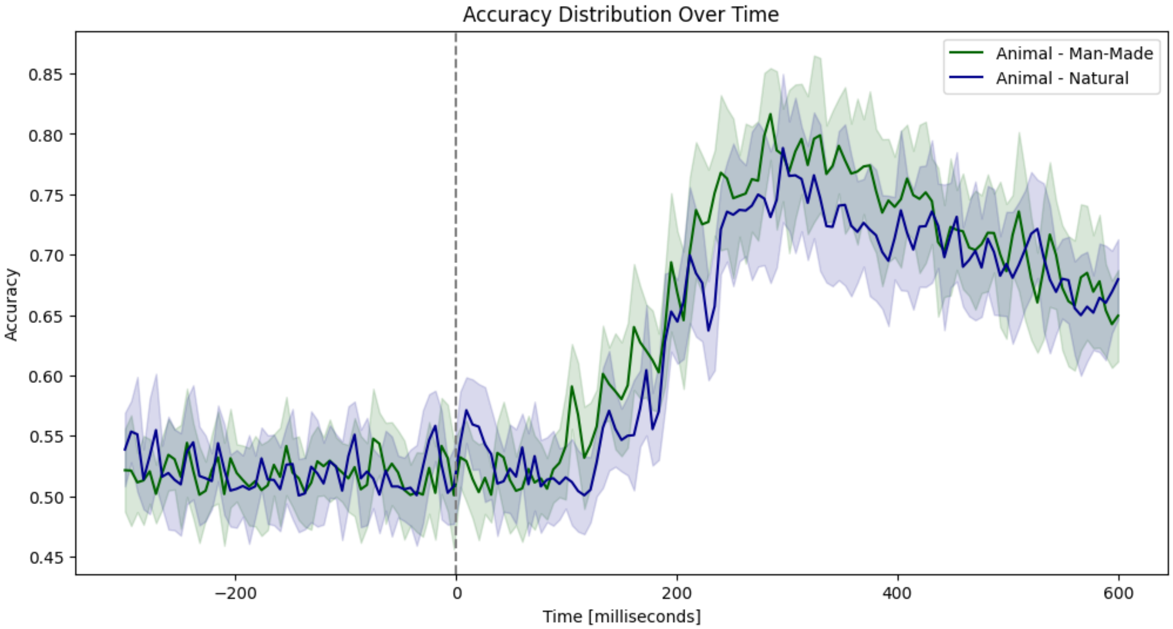


**(a)** Classification accuracy of comparing animal with man-made, and natural.

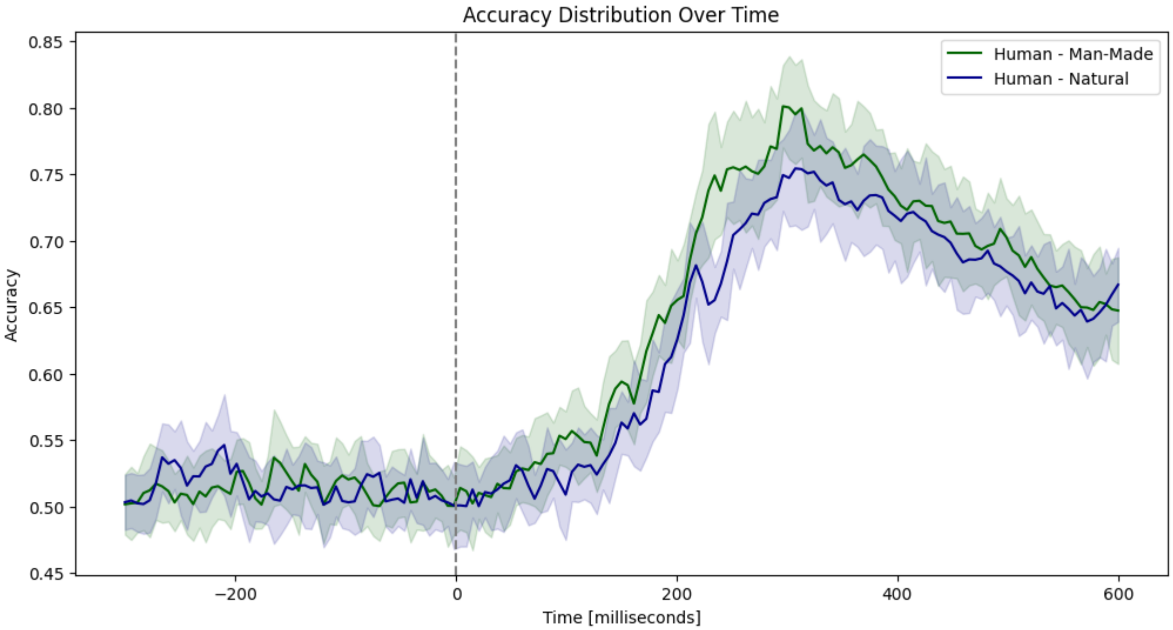


**(b)** Classification accuracy of comparing human with man-made, and natural.

**Fig. S1.** Accuracy of classifying animal, human from man-made, and natural.

Figures 3-5 (multi-category classification), the end accuracy does not converge to the chance level, indicating that some discriminative information remains. Specifically, while pairwise decoding accuracy tends to return close to the baseline accuracy at the end of the trial, multi-category decoding accuracy remains above the chance level. This section addresses these observations, provides additional analyses to clarify these trends, and eliminates any potential misinterpretations.

To ensure clarity and avoid misleading information, we conducted further analyses focusing on the accuracy of classifying specific category pairs that we have not included in the main manuscript. The pairwise comparisons analyzed are Animal vs. Man-Made, Animal vs. Natural, Human vs. Man-Made, and Human vs. Natural.

Fig. S1 illustrates the pairwise decoding accuracy for the specified comparisons, where the end accuracy does not converge to the chance level. This supports our interpretation that some pairwise discrimination remains higher than the chance level in the multi-category scenario, which is the reason for the sustained accuracy in multi-category decoding.

In multi-category decoding, the end accuracy remains above chance even though the baseline accuracy is at chance. This can be explained by aggregating information from multiple categories, providing a richer dataset for classification. The combined microsaccade patterns across different categories retain discriminative features for longer, thus maintaining accuracy above the chance level. In addition, in multi-category decoding, the classifier leverages cumulative information from all stimuli, which might include subtle but consistent patterns not evident in pairwise comparisons.

### 3. GAZE COORDINATES

To comprehensively understand the eye movement data utilized in our study, we have included the gaze coordinates of a human participant's eye movement trial in Fig. S2. The gaze coordinates represent the precise focus point on the screen at each recorded time point. As indicated in the "Methods" section, these data were collected using a high-resolution eye tracker with a sampling frequency of 500 Hz, ensuring accurate capture of rapid eye movements such as microsaccades.

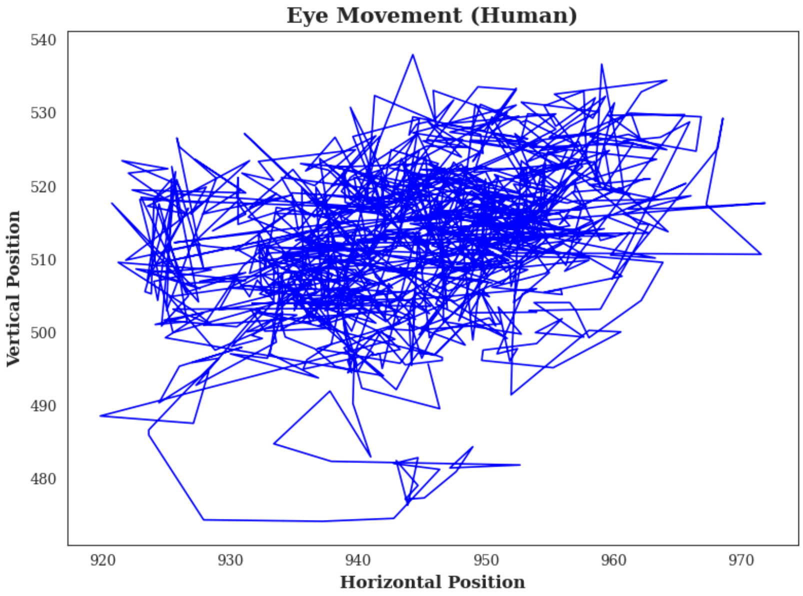


**Fig. S2.** Gaze Coordinates of human participant.

### 4. PCA OF MICROSACCADE RATE DATA

In addition to conducting pairwise comparisons using Principal Component Analysis (PCA), we also performed PCA on the entire dataset, which includes all stimulus categories. The objective was to demonstrate the variance captured by each principal component and visualize the distribution of different stimulus categories in the principal component space. We have conducted PCA on the entire dataset, including all stimulus categories, and have included the following analysis:

- **Explained Variances:** The first few principal components (PCs) and their corresponding explained variances were plotted to illustrate the variance captured by each component for each comparison and the entire dataset.

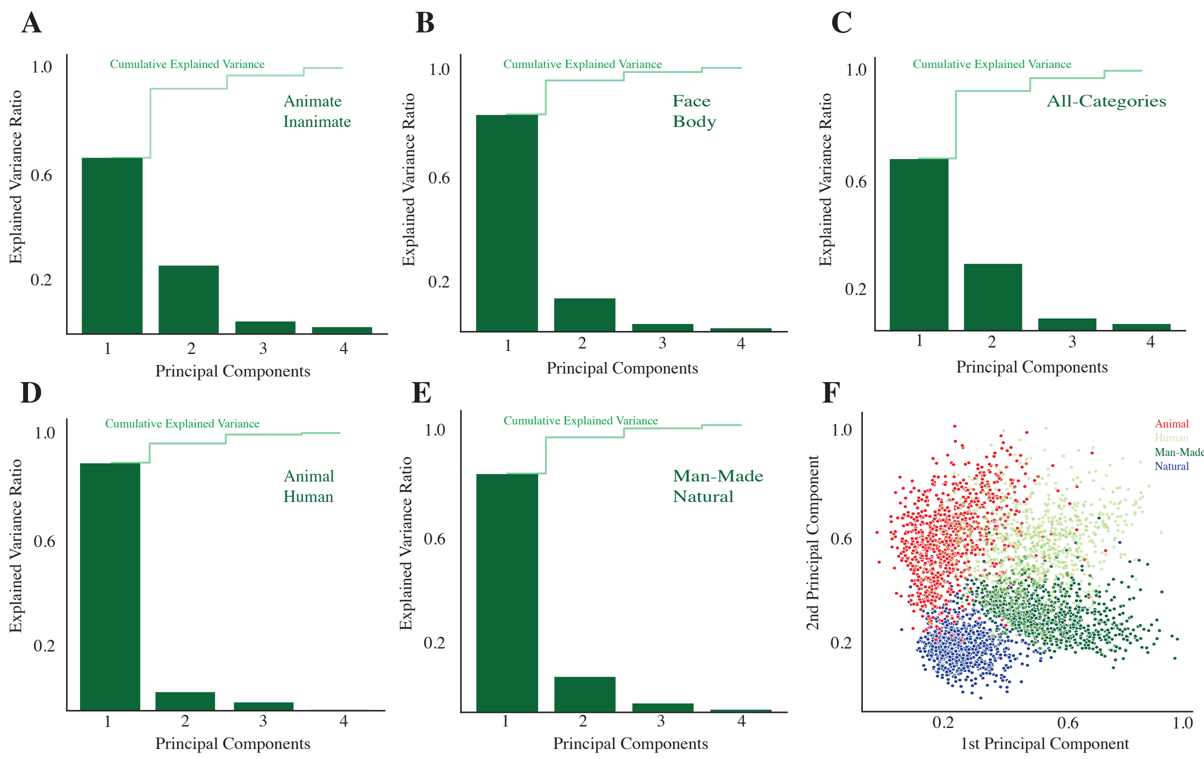


**Fig. S3.** (A-E) Cumulative explained variance for the PCs for different stimulus category comparisons. (F) 2-dimensional PCA visualization of all stimulus categories showing separability in the PCA space.

- **PCA on the Entire Dataset:** The PCA was performed on the entire dataset, including all stimulus categories.

Fig. S3(F) illustrates the 2-dimensional PCA visualization of all stimulus categories. It indicates that the distribution of stimulus categories is separable from each other. In addition, Fig. S3(A-E) represents the individual and cumulative explained variance for each pair of microsaccade rate comparisons across the stimulus categories.

The PCA analysis shows that the first four principal components capture more than 99% of the total variance in the data. The explained variance plots indicate that the first principal component alone captures most of the variance, with the first four components accounting for nearly all the variance (Fig. S3 (A-E)). The 2-dimensional PCA visualization (Fig. S3(F)) further confirms that different stimulus categories are well-separated in the PCA space. This separability highlights the distinct microsaccade patterns elicited by different stimuli, supporting the robustness of our classification results.

### 5. VALIDATION OF CLASSIFICATION PERFORMANCE WITH TRIAL-SHUFFLED DATASET

In the original classification analysis, we aimed to determine the predictive power of microsaccade rates for discriminating between stimulus categories. However, to ensure the robustness and reliability of our results, it is essential to validate the performance of our classification model against a trial-shuffled dataset. This validation step is crucial to confirm that the observed classification accuracy is not due to random chance but reflects genuine relationships between microsaccade patterns and stimulus categories.

To address this concern, we performed an analysis using a trial-shuffled dataset. The process involved randomly permuting the trial labels to disrupt any existing relationship between the microsaccade data and the stimulus categories. This method breaks the direct association between the observed microsaccades and their corresponding stimuli, serving as a control to evaluate the performance of our classification model.

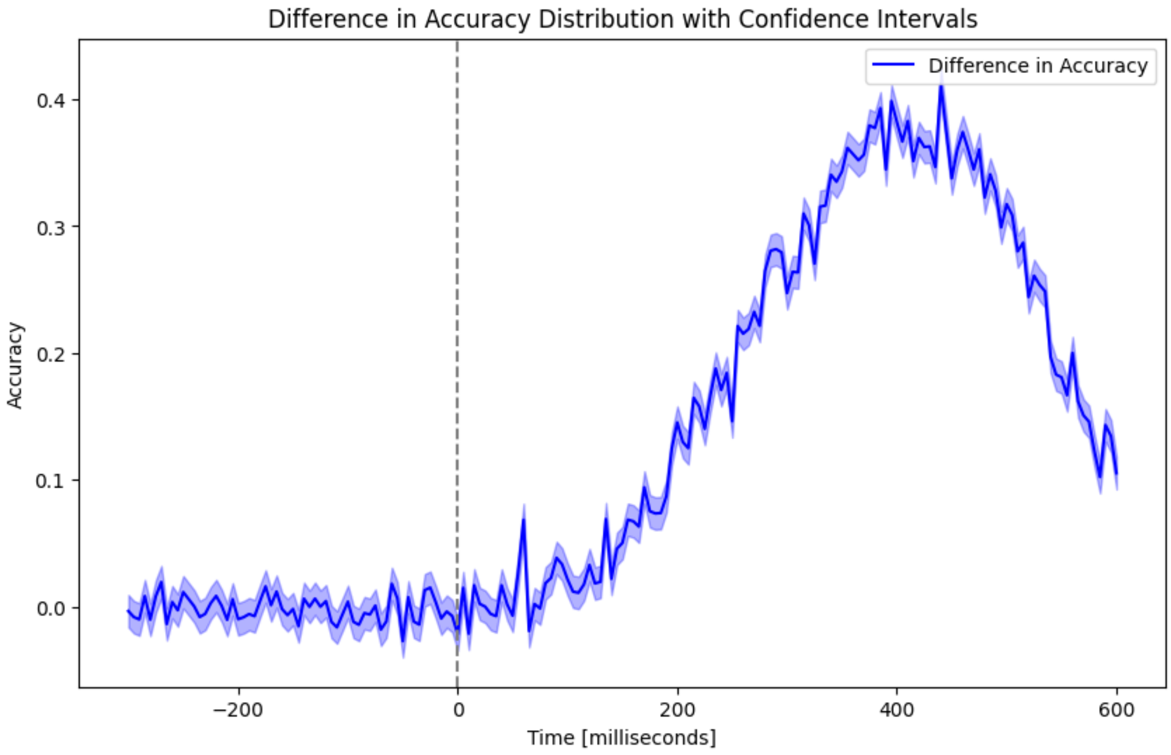


**Fig. S4.** Accuracy difference over time across classification with original dataset and trial-shuffled dataset.

We recalculated the accuracy values using the trial-shuffled dataset. As expected, the accuracy dropped significantly compared to the original dataset. This significant reduction in accuracy values for the trial-shuffled dataset indicates that the classifier's performance with the true labels is not due to random chance. Specifically, the classifier's accuracy was consistently higher for the original dataset, particularly in the post-stimulus period. The performance peaked around 400 ms after the stimulus onset and gradually decreased afterward (Fig. S4).

The comparison between the original and trial-shuffled datasets demonstrates that the observed classification accuracy values are meaningful, not data structure artifacts. The classifier's performance on the original dataset, showing significantly higher accuracy, confirms that the microsaccade rates carry discriminative information about the stimulus categories. Fig. S4 illustrates the difference in accuracy between the original dataset and the trial-shuffled dataset, highlighting the robustness of our classification model.

### 6. MICROSACCADE RATE BARPLOT

In this section, we present additional bar plots displaying the Microsaccade Rate (MSR) for different stimulus categories in both human and monkey datasets in Fig. S5. These plots complement the data presented in Figures 3B and 5B, where we reported saccade rates for different stimulus categories. It is essential to differentiate between saccades and microsaccades because they have different visual perception characteristics, frequencies, and functional roles.

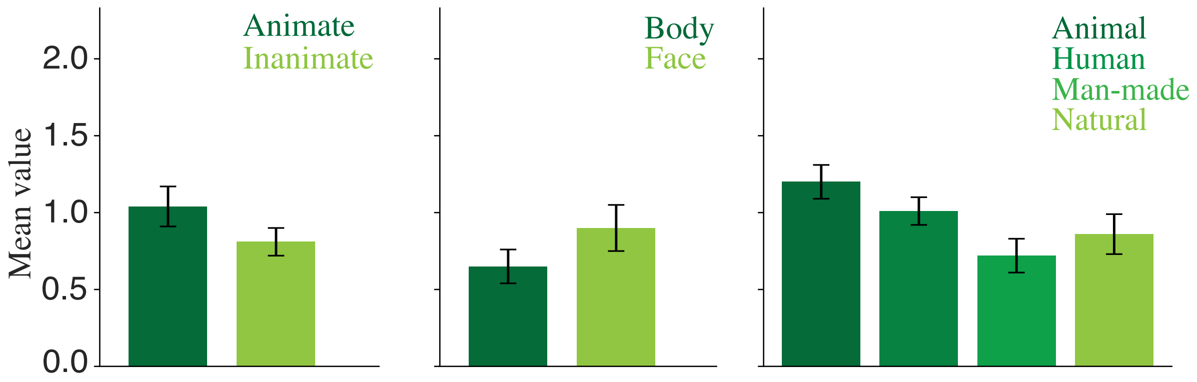


**(a)** Microsaccade rate statistics across different categories for Monkey dataset.

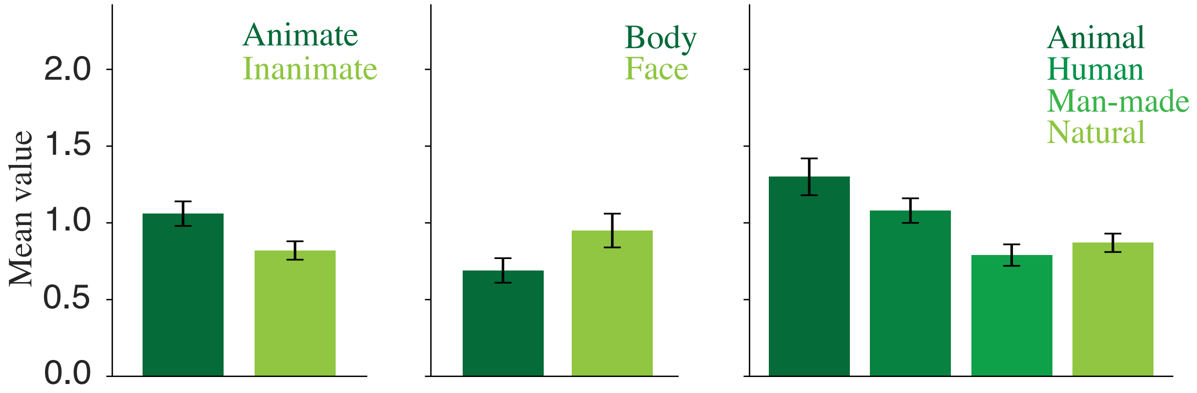


**(b)** Microsaccade rate statistics across different categories for Human dataset.

**Fig. S5.** Microsaccade rate statistics across different categories.
